# Effects of agricultural fungicide use on *Aspergillus fumigatus* abundance, antifungal susceptibility, and population structure

**DOI:** 10.1101/2020.05.26.116616

**Authors:** Amelia E. Barber, Jennifer Riedel, Tongta Sae-Ong, Kang Kang, Werner Brabetz, Gianni Panagiotou, Holger B. Deising, Oliver Kurzai

## Abstract

Antibiotic resistance is an increasing threat to human health. In the case of *Aspergillus fumigatus*, which is both an environmental saprobe and an opportunistic human fungal pathogen, resistance is suggested to arise from fungicide use in agriculture, as the azoles used for plant protection are almost identical to the frontline antifungals used clinically. However, limiting azole fungicide use on crop fields to preserve their activity for clinical use could threaten the global food supply via a reduction in yield. In this study we clarify the link between fungicide use on crop fields and resistance in a prototypical human pathogen through systematic soil sampling on farms in Germany and surveying fields before and after azole application. We observed a reduction in the abundance of *A. fumigatus* on fields following fungicide treatment in 2017—a finding that was not observed on an organic control field applying only natural plant protection agents. However, this finding was less pronounced during our 2018 sampling, indicating that the impact of fungicides on *A. fumigatus* population size is variable and influenced by additional factors. The overall resistance frequency among agricultural isolates is low, with only 1-3% of isolates from 2016-2018 displaying resistance to medical azoles. Isolates collected after the growing season and azole exposure show a subtle, but consistent decrease in susceptibility to medical and agricultural azoles. Whole genome sequencing indicates that, despite the alterations in antifungal susceptibility, fungicide application does not significantly affect the population structure and genetic diversity of *A. fumigatus* in fields. Given the low observed resistance rate among agricultural isolates, as well the lack of genomic impact following azole application, we do not find evidence that azole use on crops is significantly driving resistance in *A. fumigatus* in this context.

**IMPORTANCE:** Antibiotic resistance is an increasing threat to human health. In the case of *Aspergillus fumigatus*, which is an environmental fungus that also causes life-threatening infections in humans, antimicrobial resistance is suggested to arise from fungicide use in agriculture, as the chemicals used for plant protection are almost identical to the antifungals used clinically. However, removing azole fungicides from crop fields threatens the global food supply via a reduction in yield. In this study, we survey crop fields before and after fungicide application. We find a low overall azole resistance rate among agricultural isolates, as well a lack of genomic and population impact following fungicide application, leading us to conclude azole use on crops does not significantly contribute to resistance in *A. fumigatus*.

## INTRODUCTION

*Aspergillus fumigatus* is a globally-distributed fungus responsible for an estimated 300,000 case of invasive disease and more than 10 million cases of chronic and allergic disease globally each year (1). Humans inhale the infectious particles, or spores, on a daily basis, but are actually an accidental host for the fungus whose primary niche is soil and decaying vegetation. Management and prophylaxis against aspergillosis relies largely on the azole class of antifungals with voriconazole and isavuconazole recommended as first line therapy (2). Unfortunately, clinical resistance to the azoles in *A. fumigatus* is an increasing problem, with some medical centers reporting rates as high as 30% in specific patient populations, and similarly high rates for environmentally-isolated *A. fumigatus* (3, 4). Regrettably, the mortality for resistant infections is upwards of 90% in some patient populations (5–7). While resistance can evolve during patient therapy (8, 9), the emergence of resistance in *A. fumigatus* has mainly been linked to the use of azoles in agriculture, as structurally similar and mechanistically indistinguishable compounds are heavily used for plant protection (10). This resistance has been described as collateral damage as *A. fumigatus* is not a plant pathogen that is being directly targeted by fungicide treatments (11). The triazoles were first released for agriculture in 1973, well before they were first introduced to human medicine in the early 1990s, and are currently the most widely-utilized antifungal compound group in agriculture due to their systemic distribution in treated plants, high efficiency and broad spectrum of target pathogens (12, 13). Crops, particularly cereals and fruits, are sprayed multiple times each growing season at a recommended dose of 100 g/hectare to control powdery mildew, rust, septoria leaf blotch, and other phytopathogenic fungi (14). Currently, there are 32 azoles commercially available for plant protection (15), but only five in regular use in human medicine (16).

The most common azole resistance mechanism in *A. fumigatus* occurs via mutations in the target protein of azole fungicides, sterol 14α-demethylase (CYP51A, syn ERG11), a key enzyme of the ergosterol biosynthesis pathway. In *A. fumigatus*, the dominant resistant mechanism among both environmental and clinical isolates is a 34 bp tandem repeat (TR_34_) in the *cyp51a* promoter coupled with a leucine to histidine substitution (L98H) in the amino acid coding sequence—the net effect of which is an increase in gene expression as well as an alteration in both the stability of the target enzyme and the interaction between the protein heme cofactor of *cyp51a* and the azole ligand (17–19). Additional mutations that have been identified to confer azole resistance in *A. fumigatus* include other variations of the tandem repeat, such as TR_46_/Y121F/T289A and TR_53_, as well as other point mutations in the *cyp51a* coding sequence (20, 21).

Disease-causing fungi are responsible for roughly 20% of crop yield loss, with a further 10% loss postharvest (16), so the use of azoles is critical for securing the food supply. However, this must be balanced against the need to preserve the activity of the azoles for clinical use and, as such, there is an urgent need to identify the contributions of azole fungicide use on food crops to the development of resistance in *A. fumigatus*. We address this through systematic soil sampling conducted on 10 agricultural sites in Germany over a three-year period—including conventionally-managed fields applying azoles fungicides, as well as those practicing organic agriculture that do not use these compounds. In the largest published *A. fumigatus* sequencing effort to date, and the first to focus on the fungus in its natural niche, we also use whole genome sequencings (WGS) to examine the impact of azole fungicides on the population genetics of 64 agriculturally-isolated *A. fumigatus* isolates.

## RESULTS

### Variable abundance of *Aspergillus fumigatus* on agricultural sites in Germany

To examine the depth distribution of *A. fumigatus* in agricultural soils, we collected soil samples from a test field at five cm intervals down to a depth of 30 cm below the surface. *A. fumigatus* was most abundant in the top five cm of soil and was not significantly observed below a depth of 15 cm (Fig 1A). As a result, only the top 5 cm layer of soil was collected in subsequent soil samples.

**Figure 1.**
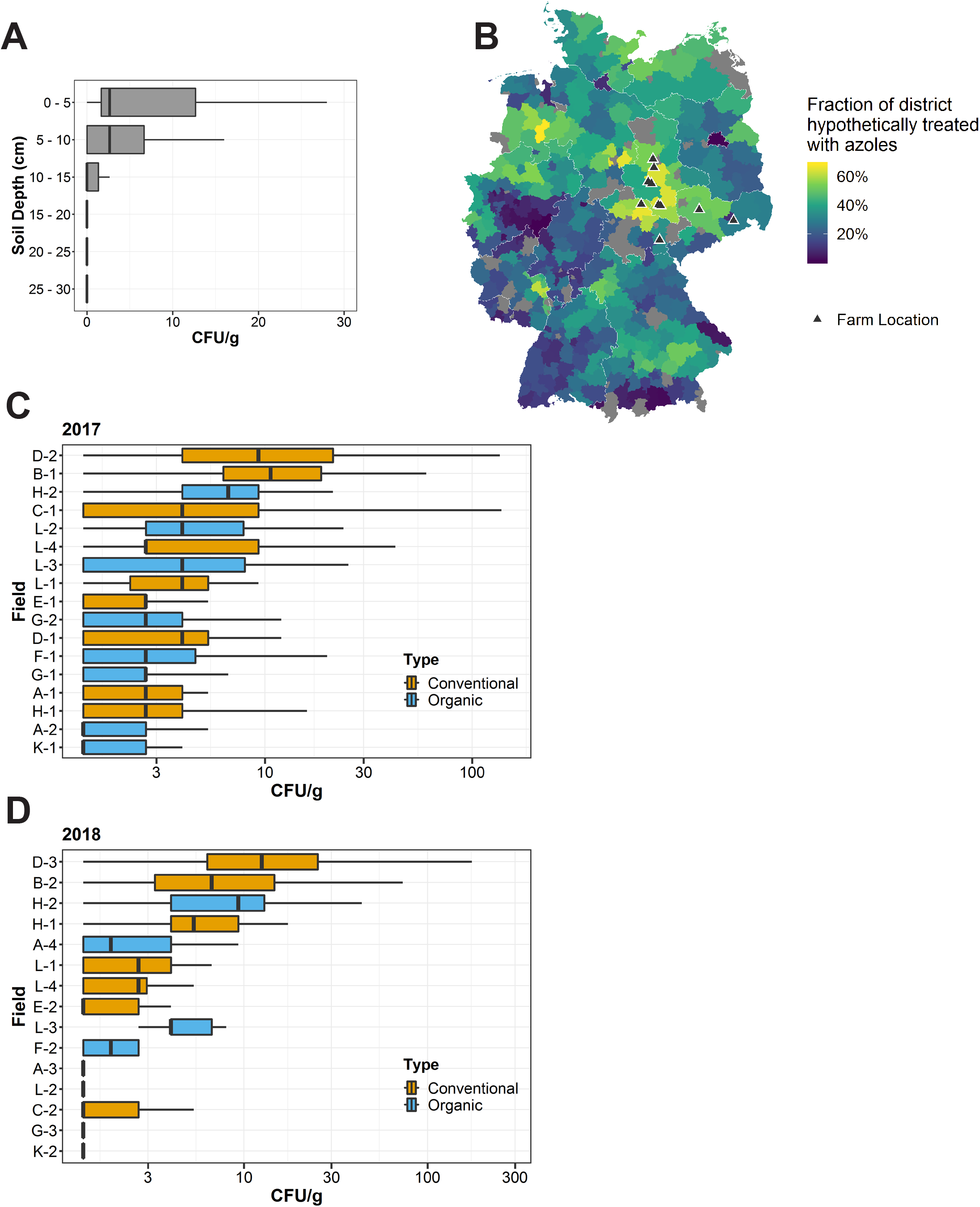
Abundance of *A. fumigatus* in the soil of conventional and organic farms. (A) CFU/g *A. fumigatus* at varying soil depths. n = 10 samples per depth. (B) Estimated fungicide treatment rates and areas in Germany. The fraction of each district that is theoretically treated with fungicides was calculated using land use and organic agriculture share data reported by the Statistical Office of Germany in December 2016. Districts where no data on land use was available are shaded grey. (C-D) Abundance of A. fumigatus in the spring as measured by CFU/g soil in the spring of 2017 (C) and 2018 (D). For 2017, boxplots represent n = 50 soil samples per field for farms A, B, C, E, F, H, K and n = 25 for farms D, G, L. For 2018, n = 50 soil samples per field for farms A, B, C, D, E, G, H, K and n = 25 for farms F and L.

To identify whether our sampling areas are representative of azole and fungicide usage for Germany as a whole, we calculated the fraction of each administrative district that is theoretically treated with azoles in the context of agriculture using publicly available data. Overall, 51% of Germany is designated as agricultural land (see Material & Methods for data sources). However, not all this land is sprayed with azoles. For example, meadow or pastureland is rarely treated with fungicides. Additionally, approximately 7% of agriculture in Germany utilizes organic farming and does not apply azoles, with a range of 3-15% between the different federal states. Using land use data on arable farmland and permanent crop areas for each district in Germany, we calculated that the mean fraction of potentially azole-treated area in Germany is 32% (for the districts where data is available), with a range of 1-68% (Fig 1B). The districts where we performed our soil sampling were almost all above the average for Germany, with a mean of 52% potentially azole-treated hectares and a range of 30-65%.

To examine the inter- and intra-field variability in the density of *A. fumigatus* on agricultural fields, soil samples were taken from nine conventional and eight organic fields before the growing season in 2017. Conventional fields were also sampled again after the vegetative period and application of azoles. In total, 2875 soil samples were taken between 2016 and 2018 (Supplemental Table 1). The predominant crops being grown were cereals such as wheat and barley, but several apple orchards were also sampled. Of the fields sampled during this period in 2017, 67% of the soil samples taken were positive for *A. fumigatus* with a large range between fields (28-100%) (Fig 1C). We also observed a large degree of variation in the mean CFU/g of soil between different fields, with some fields having 30x higher *A. fumigatus* density over others (0.7 – 18.8 CFU/g).

To examine the stability of *A. fumigatus* population sizes in agricultural soil, we investigated the same farms a year later in the spring of 2018 and repeated the soil sampling on eight conventional and seven organic fields. Due to crop rotation, it was not possible to sample the identical fields as the previous year, except for the apple orchards on Farms H and L. During the 2018 sampling period, an overall lower proportion of samples were positive for *A. fumigatus* (51% with a range of 10-96% between different fields) (Fig 1D), but the mean CFU/g for the 1000 samples was similar to the previous year (5.18 CFU/g soil for 2017 compared to 5.74 CFU/g for 2018). As in the previous year, there was a large variability in the mean CFU of *A. fumigatus* present between fields. When comparing the six apple fields that were sampled over consecutive years, we did not observe a consistent trend in the stability *A. fumigatus* population size. Two fields showed similar levels of *A. fumigatus* between 2017 and 2018, while two fields showed an increased abundance between the years, and the remaining two field showed a significant reduction in abundance (Supplemental Fig 1A). To investigate potential factors that might support a higher abundance of *A. fumigatus*, we compared the total organic carbon (TOC) content of a random selection of samples with the CFU/g *A. fumigatus,* but did not detect a clear relationship between the two (Supplemental Fig 1B). Altogether, we observed a non-uniform distribution of *A. fumigatus* in soil samples taken from the same field and a large degree of heterogeneity between fields.

### Variable effects of fungicide application on *A. fumigatus* abundance

To examine the impact of fungicide application and the azoles on *A. fumigatus* in agricultural soil, we performed additional soil sampling on the conventional fields surveyed in the spring at the end of the vegetative period and after several months of fungicidal crop protection. A schematic illustration of the soil sampling and fungicide application timelines for 2017 and 2018 can be found in Supplemental Figures 2A and B. Unfortunately, the fungicide history for Farms H and L was not available to us. Fields were typically treated with fungicides twice during the growing period and azoles were by far the most dominant class of fungicide applied. Every application recorded contained at least one azole. However, fungicides are often applied as commercially available cocktails of different chemicals, so other classes were also present in 0-55% of applications in 2017 and 2018 (summarized in Supplemental Fig 2C).

When comparing the amount of *A. fumigatus* on fields before fungicide application to after fungicide application and azole exposure, we detected a significant reduction in the CFU/g soil on the majority of fields in 2017 (Fig 2A), even though it is not being directly targeted as a plant pathogen. To investigate whether this reduction in agricultural *A. fumigatus* populations was the result of fungicide application, and not a seasonal effect from comparing April to July, monthly soil samples were taken from a conventional field and an organic field not treated with azoles or other non-natural fungicides as a control. Samples were taken beginning in April, before azoles were applied to the conventional field, through the harvest period in July, and then additional samples were taken in October and November to allow for a period without fungicide application. From April to July, the conventional field was sprayed with azoles every 3-5 weeks. When comparing the abundance of *A. fumigatus* on the organic field, we did not observe any significant differences between the abundance recorded monthly between April and July (Fig 2B). However, the conventional field showed a significant reduction in abundance between April and May, corresponding to the beginning of the azole application period, and this reduction was maintained through the rest of the azole application period (Fig 2C).

**Figure 2.**
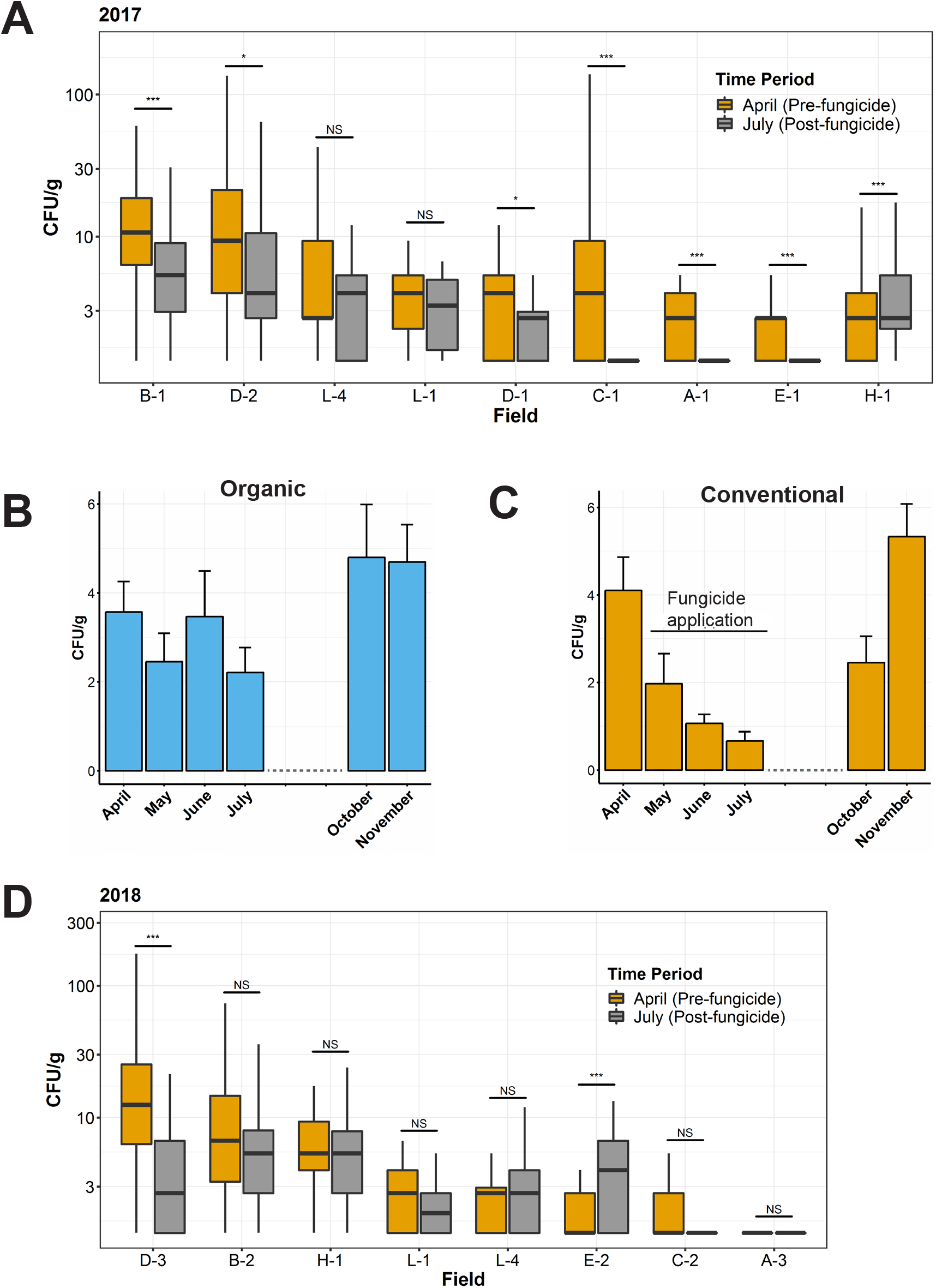
Abundance of *A. fumigatus* in the soil of conventional farms before and after the vegetative period and fungicide application. (A, D) CFU/g *A. fumigatus* on conventional fields sampled in April, prior to the vegetative period and fungicide application (orange), and in July, after the vegetative period and three months of fungicide application, including azole fungicides (gray) in 2017 (A) and 2018 (D). *, *P* s; 0.05; ** *P* s; 0.01; *** *P* s; 0.001 as determined by Mann-Whitney U test. (A) Boxplots represent n = 50 samples per field and time point for farms A, B, C, E, H and n = 25 samples for farms D and L. (D) n = 50 samples per field and time point for farms A, B, C, D, E, H and n = 25 samples for farm L. (B-C) CFU/g *A. fumigatus* during the months of April, May, June, July, October and November of a conventional field applying fungicides from May-July (C) and an organic field not applying non-natural fungicides (B). Bars represent mean ± SEM of 50 soil samples per month. No significant difference was found in abundance between the months of April, May, June, and July for the organic field using a Kruskal-Wallis test. In contrast, we found a significant difference (*P* = 0.004) for the abundances in this time period on the conventional field. Pairwise Wilcoxon signed-rank test confirmed a significant difference between April and all subsequent months until the cessation of fungicides.

When comparing *A. fumigatus* density before and after fungicide treatment in 2018, we did not observe the same reduction in abundance and most fields did not show significant changes between the time points (Fig 2D). In fact, only one of eight fields sampled showed a statistically significant reduction in *A. fumigatus* abundance. Altogether, the impact of fungicide application on *A. fumigatus* abundance was variable between fields and more so between years, as other environmental factors also appear to influence *A. fumigatus* population size in agricultural soil.

### Reduced susceptibility to agricultural azoles in populations isolated after the growing season and azole exposure

To assess the susceptibility of *A. fumigatus* to commonly-applied agricultural azoles, we screened 435 isolates from 2017 and 342 isolates from 2018 for their ability to grow at a set concentration of difenoconazole and tebuconazole (approximately 20 isolates per field and sampling point). To limit potentially clonal isolates from skewing the results, a maximum of two isolates per soil sample were included for testing. As there are no established breakpoints for defining resistance to these compounds in *A. fumigatus*, we selected concentrations that mimicked MIC90 values for these azoles (1 mg/L for difenoconazole and 2 mg/L for tebuconazole). When examining conventional and organic farms in the spring, we observed a wide range in the fraction of isolates per field that were able to grow when challenged with agricultural azoles. For difenoconazole, this ranged from 10-55% per field in 2017 (n = 17 fields, 320 isolates in total) and 0-50% in 2018 (n = 15 fields, 261 isolates in total) (Fig 3A and Supplemental Table 2). For tebuconazole, the rates ranged from 0-25% in 2017 and 0-20% in 2018 (Fig 3B and Supplemental Table 2). We did not detect any significant differences in the rates between conventional and organic fields in the spring or between fields growing different crops (cereals or apples).

**Figure 3.**
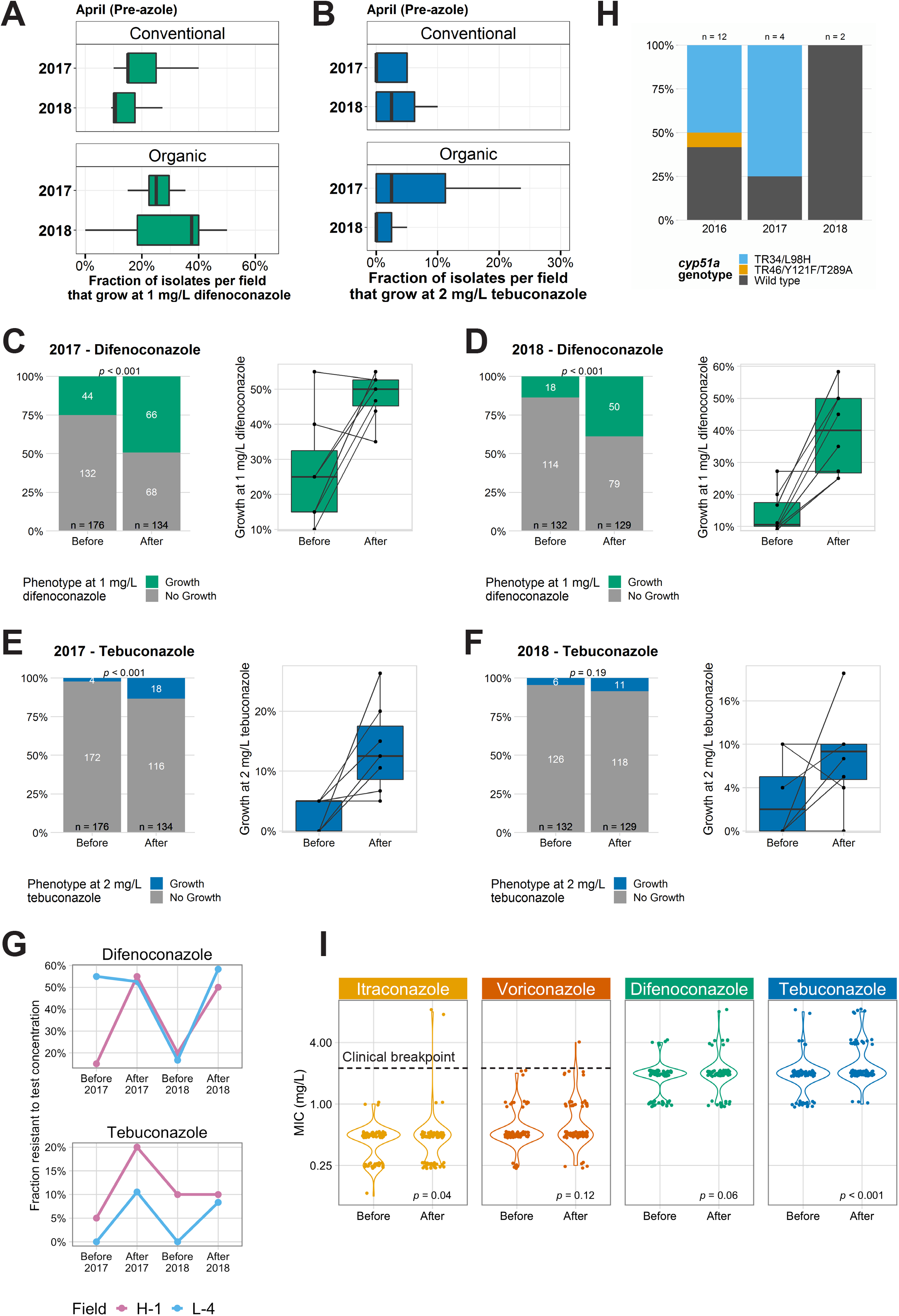
Azole resistance among agricultural *A. fumigatus.* (A-B) Fraction of the isolates per field that grow at 1 mg/L difenoconazole (A) and 2 mg/L tebuconazole (B). For 2017, n = 340 isolates from 9 conventional and 8 organic fields, ≈ 20 isolates per field. For 2018, n = 213 isolates from 8 conventional and 7 organic fields, ≈ 20 isolates per field (full summary in Supplemental Tables 2 & 3). (C-F) Comparison of the proportion of isolates that grow at 1 mg/L difenoconazole (C-D) and 2 mg/L tebuconazole (E-F) before and after the vegetative period and fungicide application. For 2017 (C, E), n = 275 isolates from 7 fields and for 2018, for 2018 n = 261 isolates from 8 fields (full summary in Supplemental Tables 4 & 5). Left panels show overall summary of all isolates tested that year. Right panels show within field changes (n ≈ 40 isolates per field; 20 before, 20 after). *P* values calculated by Wilcoxon signed-rank test between before and after values. (G) Temporal changes in antifungal susceptibility of A. fumigatus on apple fields sampled before and after fungicide exposure over a two year period. Fraction of isolates that can grow at 1 mg/L difenoconazole (top) and 2 mg/L tebuconazole (bottom). n = 12-20 isolates/field and time point. (H) cyp51a genotypes of isolates resistant to one or more medical azole. (I) MICs of agricultural *A. fumigatus* isolated before and after azole exposure. n = 159 randomly selected isolates from 2017 and 2018; n = 80 before and n = 79 after. P values calculated by Wilcoxon signed-rank test between before and after values.

Given the reduction in the *A. fumigatus* population size observed on most conventional following fungicide application in 2017, and more variably on fields in 2018, we wanted to examine the effect of fungicides on the local azole susceptibility following the vegetative period and several months of fungicide application. We observed an increase in the proportion of isolates that were able to grow at the test concentrations of difenoconazole (1 mg/L) and tebuconazole (2mg/L) for fields sampled after azole exposure compared to the same field in the spring prior to azole application (Fig 3C, D, and Supplemental Table 3). We detected a 1.97-fold increase in the proportion of isolates that were resistant to our test concentration of difenoconazole in 2017 with a range from a −1.1 to 4.5-fold for individual fields (Fig 3C). In 2018, this increased to a 2.84-fold with a range of 1-4.5-fold change increase for individual fields (Fig 3D). We also saw a similar increase in the fraction of isolates with reduced susceptibility to our test concentration of tebuconazole after azole exposure compared to the before. In 2017, we observed a 5.9-fold in number of isolates that were able to grow at the test concentration after azole exposure vs. before exposure with a range of 0-25.0% for individual fields (Fig 3E). In 2018, we detected a more modest 1.9-fold with range of −5 to 18.2% for individual fields (Fig 3F).

For the fields growing cereals, we were not able to sample the same fields over subsequent years due to crop rotation and the fields not being in use the following year. However, we were able to compare the same apple fields in both 2017 and 2018. We tracked the local susceptibility to agricultural azoles in these fields over two consecutive years at two time points: in the spring prior to fungicide application and just prior to harvest after ≈ 3 months of fungicide application. The fraction of isolates that were able to grow in the presence of 1 mg/L difenoconazole or 2 mg/L tebuconazole was, in general, low in the spring, and increased after fungicide application (Fig 3G, Supplemental Table 3). Interestingly, the spring of 2018, the proportion had returned to a level comparable to what we observed in the spring of 2017—indicating that the reduced susceptibility is transient and recedes when the selective pressure imposed by fungicide is removed. In summary, we found a wide range of susceptibilities to agricultural azoles between different fields, but a consistent decrease in susceptibility following the growing season, fungicide application, and azole exposure. However, this change is seemingly transient or reversible, and the *A. fumigatus* populations from fungicide-treated fields typically returned to what they were prior to fungicide application by the following spring.

### Resistance to medical azoles in agricultural *A. fumigatus* isolates

We next examined our isolate collection for resistance to medical azoles and determined the proportion that would be considered clinically resistant. Using the VIPcheck™ agar-based screening method followed by broth microdilution for isolates showing growth on agar-containing wells (22, 23), we determined that only a very small fraction of *A. fumigatus* isolated from agriculture showed resistance to itraconazole, voriconazole, or posaconazole in 2016-2018 (Table 1A). The overall resistance rate to itraconazole among all isolates collected was highest, with 3.0% (11/333) of isolates being resistant in 2016, 0.7% in 2017 (4/460), and 0.6% (2/322) in 2018. We observed lower resistance rates for posaconazole and voriconazole, with only 2.1% (7/333) being resistant in 2016, 0.7% (4/460) in 2017, and 0.0% (0/322) in 2018. As there are no clinical breakpoints established for agricultural azoles, we calculated epidemiological cutoff values (ECOFFs) for these difenoconazole and tebuconazole using MICs from 160 randomly selected isolates from 2017 and 2018. Using these values, we found that 1.3% of the 160 isolates had MICs above the ECOFF for difenoconazole (2 mg/L) and 4.4% of isolates had MICs above the ECOFF for tebuconazole (2 mg/L) (Table 1B). As has been described previously (24), isolates resistant to one or more medical azoles, often displayed elevated MICs to agricultural azoles, indicating cross resistance (Supplemental Table 4).

**Table 1.** Azole resistance rates among agriculturally isolated *A. fumigatus* isolates. (A) Isolates were screened for potential azole resistance to itraconazole (ITR), posaconazole (POS), and voriconazole (VOR) using the VIPcheck agar-based screening. Resistance was confirmed and MICs determined via EUCAST broth microdilution testing. (B) MIC_50_ for medical and agricultural azoles calculated from 160 randomly selected isolates, as well as ECOFF_95_ and the fraction of isolates with MICs above this value.

| Year | n | Resistance rate (%) |  |  |
| --- | --- | --- | --- | --- |
|  |  | ITR | POS | VOR |
| 2016 | 333 | 3.0 | 2.1 | 2.1 |
| 2017 | 460 | 0.7 | 0.7 | 0.7 |
| 2018 | 322 | 0.6 | 0 | 0 |

(B)
| Azole | MIC<sub>50</sub><sup>+</sup> | ECOFF* | Fraction of isolates above ECOFF (%) |
| --- | --- | --- | --- |
| Itraconazole | 0.5 | 2 | 1.3 |
| Voriconazole | 0.5 | 2 | 0.6 |
| Difenoconazole | 2 | 8 | 1.3 |
| Tebuconazole | 2 | 8 | 4.4 |

<sup>+</sup>Calculated from 160 randomly selected isolates
\*Rounded up to next dilution

To quantify what mutations were responsible for azole resistance in the population analyzed, we genotyped the *cyp51a* locus of all isolates resistant to one more medical azole using Sanger sequencing (n = 18). This gene is the target of the azoles and the major source of resistance among both environmental and clinical *A. fumigatus*. We found that the most dominant DNA alteration observed was the well-characterized TR_34_/L98H mutation (Fig 3H, Supplemental Table 4). This *cyp51a* genotype accounted for 6/12 (50%) of resistant isolates in 2016 and 3/4 (75%) in 2017. In 2018, we identified only two resistant isolates among the 322 screened and both had wild type *cyp51a* loci. However, both of these isolates were only weakly resistant to itraconazole, but not other azoles, with itraconazole MIC values right at the breakpoint of 2-4 mg/L. For comparison, resistant isolates from other years had MIC values of >8 mg/L. In total, the genetic cause of resistance remained unknown for 8 isolates from 2016-18.

Since the majority of fields had no resistant isolates, it was not possible to effectively compare resistance rates among agricultural *A. fumigatus* for medical azoles before and after fungicide treatment. In lieu of this, we examined the MIC distribution for isolates collected before and after the growing season and fungicide application. Examining 79 isolates from the before period and 79 isolates from after azole exposure, we observed a shift in the MIC distribution towards higher MICs for all azoles examined, both medical and agricultural (Fig 3G). However, the median MICs remained unchanged, indicating that the majority of the population following azole exposure does not exhibit a change in MIC. Taken together, these results indicate that the rate of resistance to clinical azoles among environmental *A. fumigatus* isolated from agricultural environments is overall low and that exposure to agricultural azoles causes a minor increase in MIC values for some of the population, but the majority of isolates are left unchanged.

### No population structure observed for *A. fumigatus* in agricultural fields

To better understand the population structure of *A. fumigatus* in the environment and how it is impacted by the azoles, we performed whole genome sequencing on isolates from four conventional farms collected before and after azole exposure. 64 isolates were sequenced by Illumina paired-end sequencing, representing eight isolates per farm and time point. Isolates from each field and time point were randomly selected, with a maximum of two per soil sample to avoid sequencing of clonal isolates. Raw reads were checked for quality and then aligned to the Af293 reference genome, resulting in a median depth of coverage after mapping of 31.5X (range 10-90X) and a median genome coverage median of 94% (range 92.8-96.7%). Consistent with what has been observed among sequenced clinical strains (25), isolates differed from the Af293 reference by a median of 84,690 single nucleotide variants (SNVs) or 2.88 SNVs/kb, with a range of 65,854-146,055 SNVs. The analysis of copy-number variations (CNVs) identified 8,277 unique CNVs in total, with a median of 3,115 CNVs per isolate (range 1,666-4,532 CNVs). These CNVs were further delimited into a median of 2,589 deletions (range 1,247-4,405) and 5,687 insertions (range 3,872-7,030).

A maximum-likelihood phylogeny based on SNVs indicated no population stratification among isolates from different farms and regions of Germany (Fig 4). To more directly assess the association between genetic and geographic distance, we performed a Mantel test correlating a geographic distance matrix with the F_ST_ genetic distance matrix, and observed no significant association between the two. When considering the azole-resistance status of the isolates, the two itraconazole-resistant TR_34_/L98H isolates sequenced clustered next to each other, despite originating from separate farms, while the third itraconazole-resistant isolate with an undefined resistance mechanism was on a distinct branch. Despite being the nearest sequenced neighbors, the two TR_34_/L98H isolates were genetically distinct, each possessing 19,439 and 60,841 unique SNVs not shared by the other isolate, along with 67,196 common SNVs relative to Af293.

**Figure 4.**
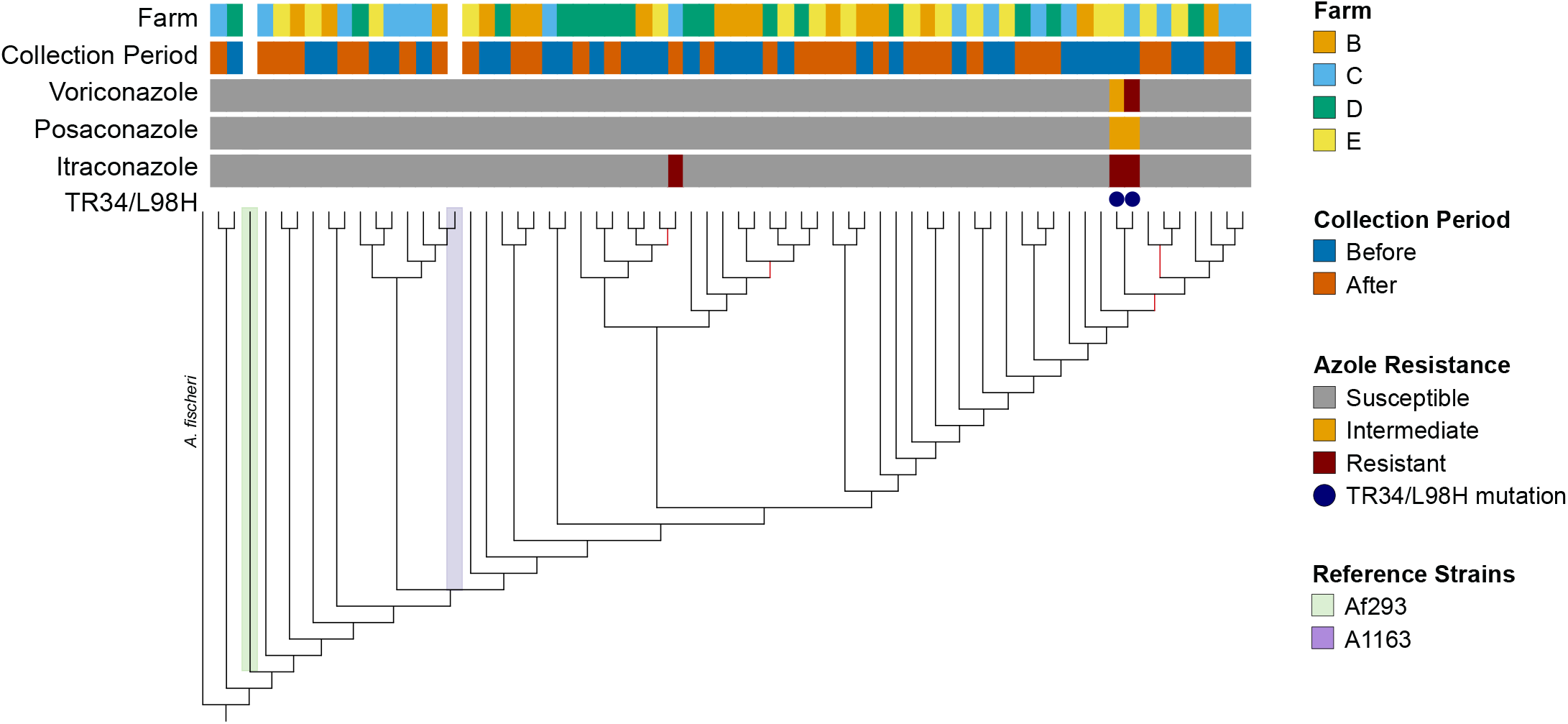
Phylogeny of agricultural *A. fumigatus* isolates from before and after the vegetative period and fungicide application. From top to bottom, the colored bars indicate: 1) the farm where the isolate was collected, 2) the collection period, 3) voriconazole resistance (susceptible, intermediate, or resistant, according to EUCAST definitions), 4) posaconazole resistance, 5) itraconazole resistance, and 6) the presence of the *cyp51a* TR34/L98H allele. *A ischerei* is indicated as an out group and the two *A. fumigatus* reference strains, Af293 and A1163 (CEA10), are also marked. Branches with support values less than 0.9 are marked in red.

Comparative analysis of molecular variance (AMOVA) indicated that the majority of the variation seen among the 64 isolates came from the population as a whole (within sample) (94.8%) and between samples (5.0%) (Table 2). There was no significant molecular variance between farms (0.2%), with the exception of modest variation between Farm B and Farm C (1.2% of variation observed). Weighted Weir and Cockerham's fixation indexes (F_ST_) for each farm were essentially zero, indicating an interbreeding, panmimetic population with no separation between farms (Supplemental Fig 3A). V_ST_ values estimating population differentiation based on copy number variation also indicated no subdivision among the farms (Supplemental Fig 3B). To examine the genetic diversity within farms, nucleotide diversity (the average number of nucleotide differences per site for all possible pairs in the population or *π)*, and the number of polymorphic sites (Watterson estimator or θ) were calculated along 5 kb windows with a 500 bp step size for each farm population. Farm C showed the greatest intra-farm diversity, while Farm E showed the smallest (Supplemental Fig 3C-D). Taken together, there was no population differentiation between *A. fumigatus* isolates from different farms in Germany, a finding in line with the fungus' capacity for aerosol dispersal.

**Table 2.** Analysis of molecular variance (AMOVA) between farms. Calculated using isolates from before and after azole application. n = 16 isolates per farm.

|  | Total variation | Variation within samples (%) | p-value | Variation between samples (%) | p-value | Variation between farms (%) | p-value |
| --- | --- | --- | --- | --- | --- | --- | --- |
| All farms | 289735.7 | 274622.7 (94.8) | <0.001 | 14629 (5) | <0.001 | 484 (0.2) | 0.303 |
| B vs. C | 147358.0 | 139257.7 (94.5) | <0.001 | 6386.2 (4.3) | <0.001 | 1714.1 (1.2) | 0.028 |
| B vs. D | 190425.2 | 181470 (95.3) | <0.001 | 9002.4 (4.7) | <0.001 | -47.2 (0.0) | 0.429 |
| B vs. E | 162874.7 | 155745.5 (95.6) | <0.001 | 6879.3 (4.2) | <0.001 | 249.9 (0.2) | 0.211 |
| C vs. D | 130625.7 | 122692.8 (93.9) | <0.001 | 7932 (6.1) | <0.001 | 1 (0.0) | 0.355 |
| C vs. E | 105529.1 | 99521.4 (94.3) | <0.001 | 6098.3 (5.8) | <0.001 | -90.6 (-0.1) | 0.414 |
| D vs. E | 143684.1 | 135847.0 (94.5) | <0.001 | 8382.7 (5.8) | <0.001 | -545.6 (0.4) | 0.890 |

### Changes in population genetics following azole exposure vary by field

Given the observed reduction in overall *A. fumigatus* abundance after azole treatment, we examined the populations for changes in genetic diversity and evidence of selective sweeps in *A. fumigatus* field populations. Using 5 kb sliding windows with a 500 bp step size, nucleotide diversity (*π*) was calculated for the individual farms before and after azole treatment. No clear trend was seen regarding changes in nucleotide diversity between the time points. The isolates from Farms B and E showed increased diversity following azole application, while nucleotide diversity decreased on Farms C and D (Fig 5A). We also calculated the number of segregating sites (*θ*) for the same 5 kb windows and observed the same lack of consensus. Farms B and C showed similar values of *θ* before as compared to after azole application, while Farm D showed a dramatic decrease in *θ* following azole application (Fig 5B). Molecular variance analysis (AMOVA) on isolates from before and after the vegetative period and azole exposure did not indicate any significant molecular variation between the time periods, with the majority of the variation being between and within samples (Table 3). Finally, we measured Tajima's D to test neutrality along 5 kb windows, where negative values indicate less variation than expected and are indicative of a selective sweep. Positive values denote a population that is more heterogenous than would be expected and suggest either a sudden population contraction or balancing selection. Overall, the bulk of the Tamija's D values were close to neutral and there was no clear trend between farms, indicating that there was no genomic signature of a population bottleneck or selective sweep following azole exposure (Fig 5C). The median Tajima's D was roughly zero for Farm B and increased slightly to 0.37 following azole exposure prior to azole application, and the same direction shift was seen for Farm D, but the starting Tajima's D was negative at the time point before azole exposure (−0.53 to 0.53) (Fig 5C). Conversely, the median Tajima's D for farms C and E shifted from positive to negative after the growing season and azole exposure (0.75 to −0.57 for field C and 0.61 to −0.10 for Field E). Taken together, these results indicate that despite the reduction in abundance of *A. fumigatus* on agricultural fields following azole application, we were unable to detect any marked changes at the population level.

**Table 3.**
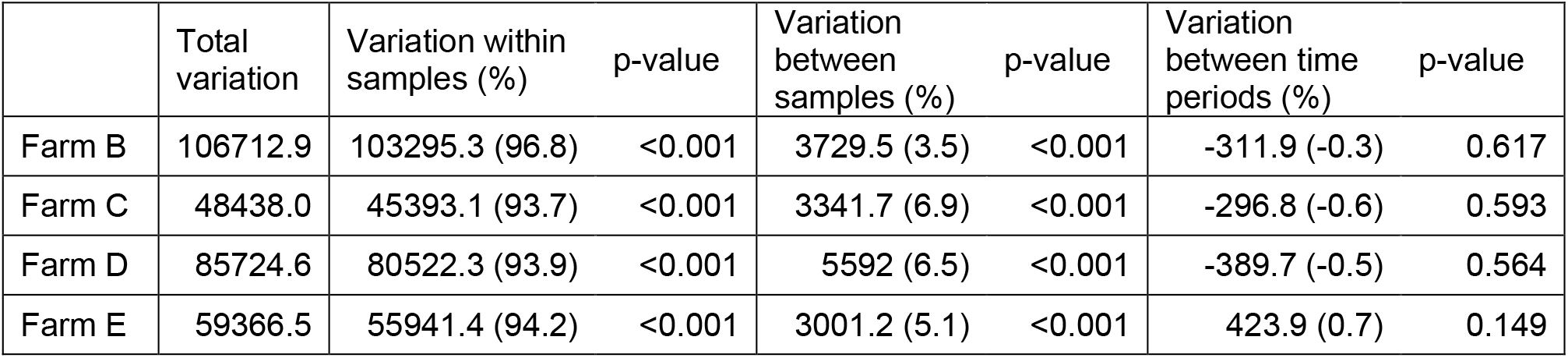
Analysis of molecular variance between isolates collected before and after azole exposure. n = 16 isolates per farm field (eight pre-azole and eight post-azole).

|  | Total variation | Variation within samples (%) | p-value | Variation between samples (%) | p-value | Variation between time periods (%) | p-value |
| --- | --- | --- | --- | --- | --- | --- | --- |
| Farm B | 106712.9 | 103295.3 (96.8) | <0.001 | 3729.5 (3.5) | <0.001 | -311.9 (-0.3) | 0.617 |
| Farm C | 48438.0 | 45393.1 (93.7) | <0.001 | 3341.7 (6.9) | <0.001 | -296.8 (-0.6) | 0.593 |
| Farm D | 85724.6 | 80522.3 (93.9) | <0.001 | 5592 (6.5) | <0.001 | -389.7 (-0.5) | 0.564 |
| Farm E | 59366.5 | 55941.4 (94.2) | <0.001 | 3001.2 (5.1) | <0.001 | 423.9 (0.7) | 0.149 |

**Figure 5.**
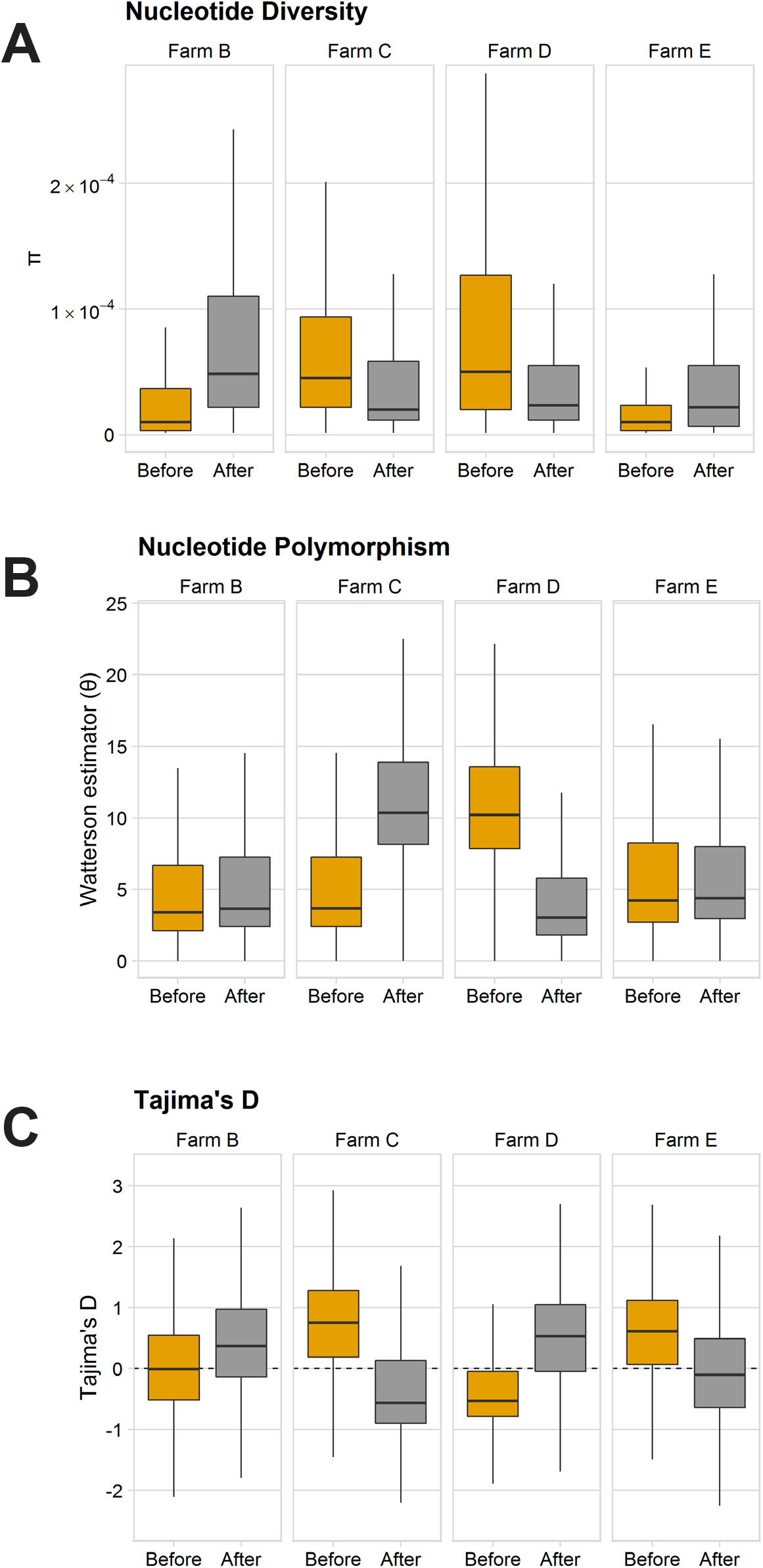
Genetic diversity among isolates from before and after the vegetative period and fungicide application, including azole exposure from four farms. (A-C) Nucleotide diversity (π) (A), nucleotide polymorphism (Watterson estimator or θ) (B), and Tajima's D (C) along 5 kb windows before and after the vegetative period and fungicide application, including azole exposure. n = 8 isolates per farm and time point, 64 in total.

## DISCUSSION

The use of azole fungicides for plant protection has been previously suggested as a driver of clinical resistance in the environmental saprobe and human pathogen *A. fumigatus*, as well as the emergence of new fungal pathogens such as *Candida auris* (11, 26, 27). However, direct evidence linking azole use in agriculture and clinical resistance is missing. Additionally, delineation of specific roles for the use of azoles in crops vs. ornamental plants such as flower bulbs has not been defined and is important to make, as limiting azole fungicide use would significantly impact disease control and yield in many crops. In this study, we provide the first systematic investigation of the impact of fungicide use on the ecology and azole resistance status of a human pathogen, *A. fumigatus*. Through analysis of 2,875 soil samples over a three-year period in central Germany, we found an overall low incidence of *A. fumigatus* isolates that would be considered clinically resistant (1-3%). However, we observed a modest, but consistent decrease in azole susceptibility following the growing season and azole exposure, as well as a more variable reduction in fungal abundance following fungicide application. Interestingly, this change in susceptibility was transient and reset by the following spring. We also assessed the influence of fungicide application on *A. fumigatus* population dynamics by WGS and were unable to find a clear impact on the population structure or genetic diversity.

Despite sampling on fields that were actively treated with azole fungicides, and in regions of Germany with above average fungicide exposure, only 1-3% of *A. fumigatus* isolates collected were resistant to medical azoles. This incidence is in agreement with clinically-reported frequencies in Germany and other European countries, where the rate of triazole resistance ranged from 0.6% to 4.2% (3.2% in Germany) (28, 29). The environmental resistance rates reported for Europe, however, have been much broader. Some studies have found environmental resistance frequencies approaching 20%, while others have reported virtually no incidence of resistant environmental isolates (30–35). Part of these differences could be attributed to differences in methodology, such as the usage of azole selection during the isolation procedure or inclusion of potentially clonal isolates from a given sample to skew the data. In the current study, we avoided azole selection during the isolation procedure, as well as only allowed two isolates per soil sample, to avoid potentially clonal isolates that influence the results. Another potentially contributing factor could be that while all technically “environmental” isolates, rural or agricultural settings could be a completely different niche than an urban flower garden—a supposition supported by a recent study where rural areas in the UK had much lower resistance rates (1.1%) than urban locations (13.8%) (34).

Another important and novel finding from this study is that isolates from after the vegetative period and azole exposure consistently showed decreased azole susceptibilities to difenoconazole and tebuconazole, as well as the subtle MIC shift to both agricultural and medical azoles. However, one shortcoming to our study is that we did not collect isolates from organic fields at a matching time point for susceptibility testing, so we cannot exclude that the changes observed in azole susceptibility are not also influenced by seasonal changes. The observation that changes in susceptibility are transient and reset in the period between the end of the growth period and the following spring on the two field is intriguing and worthy of further study. This transformation could either be the result of a naïve population coming in via aerosol dispersal, or a consequence of isolates acquiring unstable, epigenetic-mediated resistance—a phenomena previously observed in the environmental saprobe and human pathogen *Mucor circinelloides* as well as the plant-pathogenic fungi *Monilinia fructicola* following azole exposure (36, 37). Either scenario would be in agreement with our finding that fungicide application does not alter the population structure of genetic diversity of *A. fumigatus* in agricultural fields.

We observed a wide range in the CFU/g soil within and between farms during annual spring sampling. Our mean *A. fumigatus* CFU/g soil was in line with what was reported recently for abundance in wheat grain, maize silage, and fruit waste (38). However, this abundance is several magnitudes lower than that reported for *A. fumigatus* in flower bulb waste and green material waste in this same study, where it was not uncommon to isolate 10^4^ CFU/g. We demonstrated a reduction in the *A. fumigatus* population size on most fields sampled in 2017 following the vegetative period and fungicide application. However, this finding was not strongly observed in 2018, indicating that other environmental factors also influence the abundance of *A. fumigatus* in agricultural soil. One example of such a potential factor is that Germany experienced extreme heat and drought during the 2018 growing season and, in fact, one field had to be removed from analysis this year because it caught fire during the vegetative period.

Our study also provides the first WGS-based study focused on *A. fumigatus* in its natural niche, the environment. Previous studies have primarily concentrated on clinical isolates, with particular priority given to resistant strains (25, 39). Even while sampling within the same field, we found a large degree of genetic diversity, where the majority of diversity came from within samples. We also did not observe a defined population structure or separation between farms or regions. The degree to which this environmental diversity is recapitulated in clinical isolates, and whether there are enrichments for particular subgroups in the transition from environment to clinic is an interesting question for further study.

Given our low observed resistance rate among agricultural isolates, as well the lack of genomic impact of azole application on the fields surveyed, our study does not find evidence that azole fungicide use in crop agriculture significantly contributes to resistance in *A. fumigatus,* nor should be necessarily removed from use in this context due to their crucial role in global food production. Instead, we propose a refined model, where azole resistance in *A. fumigatus* is being driven by the use of these compounds in the cultivation of flowers and other ornamentals. It is common practice to dip flower bulbs directly in fungicide stocks prior to storage and shipment, and their use in this setting should be more carefully considered and managed (40, 41). This revised model is supported by the recent identification of flower bulb and green waste as environmental “hotspots” for resistance selection, which possess both high overall fungal colony counts as well as increased incidence of resistant *A. fumigatus* isolates compared to soil (38). Unfortunately, due to massive aerial dispersal of *A. fumigatus* conidia, the use of azoles in any hotspot can lead to the worldwide distribution of resistant strains.

## MATERIALS AND METHODS

### Site selection and soil sampling

During 2016 to 2018, soil sampling was conducted on agricultural sites in the federal states of Thuringia, Saxony-Anhalt, and Saxony with the approval of the land owner and/or relevant ministries. The majority of fields sampled were growing cereals such as wheat or barley, but some apple orchards were sampled as well (see Supplemental Table 1 for full details). Farms were arbitrarily assigned an alphabetic identifier (A-L) and specific fields a numeric identifier as described in the supplementary Materials & Methods. Due to crop rotation, the same field could not be surveyed over subsequent years, with the exception of the apple orchards. Soil samples were collected at beginning of the vegetation period on the conventional and organic cultivated sites and after azole application an additional sampling was carried out on the conventional sites. In general, 50 soil samples per site and type of farming (conventional or organic) were collected, with a total of approximately 1000 soil samples per year. Soil samples were selected to best cover the field with a minimum distance of 1 m between samples. For each sample, the top layer of soil was collected by a metal spatula into a sterile sample cup and refrigerated until processing.

### Soil processing and isolation of *A. fumigatus*

3 g of soil from each sample cup was weighed out and resuspended in 8 mL 0.2M NaCl containing 1% Tween20. Samples were vortexed vigorously and then left to settle until a phase separation became apparent. 2 mL of the upper phase was transferred to a new tube for plating onto Sabouraud Glucose Agar (SGA) containing 50 μg/mL Chloramphenicol (Sigma-Aldrich, Taufkirchen, Germany). Of this 2 mL, 150 μL was plated onto one plate and the remaining volume was plated onto a second plate in order to adjust for variable fungal concentrations in samples. Plates were then incubated at 50°C for 5 days to select for *A. fumigatus,* which is unique among *Aspergillus* spp. in its ability to grow at this temperature. On day 5, the incubator temperature was reduced to 42°C to allow for sporulation and plates grown for another 2 days. The number of *A. fumigatus* colonies was counted and up to two colonies per soil sample transferred to new plates for isolation.

### Antifungal susceptibility testing

Quick screening of susceptibility to itraconazole, voriconazole, and posaconazole were assessed using the agar-based VIPcheck™ Assay (Mediaproducts BV, Groningen, Netherlands) following the manufacturer's directions. For testing, isolates were grown on SGA for 2-4 days at 37°C. Plates were then swabbed with a damp, sterile cotton swab to prepare a conidial suspension of 0.5-2 McFarland. 25 μL of this suspension was then plated onto the 4 wells of the VIPcheck plate containing 4 mg/L itraconazole, 2 mg/L voriconazole, 0.5 mg/L posaconazole, or a control well containing no drug. To assess susceptibility to agricultural azoles during isolates collected during 2017 and 2018, RPMI+2% glucose agar plates containing either 1 mg/L difenoconazole or 2 mg/L tebuconazole were prepared as in (42). Resistance was defined as significant inhibition of germination and hyphal growth compared to the no drug control. Isolates that showed resistance to any of the medical azoles in the VIPcheck assay were subject to broth microdilution following EUCAST methodology (protocol E.DEF 9.3). ECOFFs were calculated using the ECOFFinder program available from EUCAST.

### DNA extraction

For WGS and PCR-based amplification, isolates were grown shaking in SG broth at 37°C and genomic DNA isolated using the Quick-DNA Fungal/Bacterial miniprep kit (Zymo Research, Irvine, CA) according to the manufacturer's suggested protocol.

### *cyp51a* genotyping

The *cyp51a* coding sequence and upstream region containing the tandem repeat was amplified using the primers described in Supplemental Table 5. Cleaned up PCR products were sequenced and the *cyp51a* genotype determined using FunResDB (https://elbe.hki-jena.de/FunResDb/index.php).

### Genome sequencing, quality assessment, and alignment

Library preparation and 2×150bp paired-end sequencing was performed on a NextSeq 500 v2 by LGC Genomics (Berlin, Germany) following the manufacturer's recommended protocols. Sequence data quality control and filtering were performed using an in-house script and FastQC (v0.11.5). Quality reads were mapped to the *A. fumigatus* Af293 reference genome (version 2015-09-27; retrieved from FungiDB (43)) using BWA-MEM (version 0.7.8-r779-dirty) (44). PCR duplicates were marked using MarkDuplicate from Picard version 2.18.25 embedded in the Genome Analysis Toolkit (GATK; version 4.1.0.0). All WGS samples included for analysis possessed greater than 10 fold genome coverage after mapping and more than 90% of reads mapped to the reference genome.

### Variant identification and SNV-based phylogeny

Short variants, including single nucleotide variants (SNVs) and short insertions and deletions (InDels) were detected using GATK Haplotype Caller following their recommended best practices for single calling (45). Copy number variants (CNVs) were identified using Control-FREEC (46). For the phylogenetic analysis, nucleotide consensus sequences were extracted from vcf files using VCFtools (47) and an in-house script used to translate nucleotide to protein-coding sequences. Multiple sequence alignment was performed using MUSCLE v3.8.31 (48) using 7,771 conserved core genes and an approximately-maximum-likelihood phylogeny constructed using FastTree2 (version 2.1.10) (49). Interactive Tree of Life (iTOL) v4 was used for visualization (50).

### Genetic diversity analyses

Analysis of molecular variance (AMOVA) was determined using the R package ade4 (nrep = 999). Nucleotide diversity (T) was calculated by VCFtools (version 0.1.6) using 5kbp windows with a step size of 500bp. Nucleotide polymorphism (8) and Tajima's D were calculated using ANGSD (51) using a window size of 5 kbp and a step size of 500 bp. Weighted Weir and Cockerham's F_ST_ values were calculated using VCFtools, while V_ST_ values were calculated as in (52). A Mantel test correlating geographic distance matrices with pairwise FST matrices was performed using the R package ade4 (nrep = 9999).

### Estimation of fungicide treatment areas and rates in Germany

The fraction of each district theoretically treated with fungicides was calculated using publicly reported data from 2016 available from the German Statistical Offices (statistikportal.de) using the following equation:

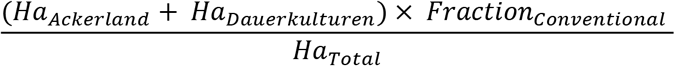

The total area per district was obtained from Table 33111-01-02-4: Ground area by actual use. The sum of arable farmland and permeant crop areas was calculated for each administrative district using Table 41141-01-01-4: Farms and their agricultural used area by crop type. To accommodate that some percentage of this area is cultivated under organic agriculture methods, the hectares of crop land was then multiplied by the fraction of non-organic agriculture for the federal state in which the district is located to estimate the number of hectares potentially treated with fungicides (as calculated using data on "Farms, agricultural areas, and workers"; accessed on November 1, 2019 on https://www.statistikportal.de/node/254). Unfortunately, no data was available on the breakdown of agricultural methods at the district level to allow for more exact estimation. Finally, this value of estimated treated area per district was divided by the total area of the district for visualization as a choropleth map.

### Box and whisker plots

Box and whisker plots presented in this paper are in the style of Tukey, where the bold line indicates the 50^th^ percentile, and the hinges represent the 25^th^ and 75^th^ percentiles. The lower whisker extends from the lower hinge to the lowest datum within 1.5 interquartile range (IQR), while the upper whisker represents the highest datum still within 1.5 IQR. Outliers are marked with points.

### Data and isolate availability

Isolates generated within this study were submitted to and are publicly available in the Jena Microbial Resource Collection. Raw FASTQ files were uploaded to the NCBI Sequence Read Archive and are publicly available under BioProject PRJNA595552.

## ACKNOWLEDGEMENTS

The authors thank Ivo Schliebner for helpful scientific discussions and his input establishing the network of farms sampled in this study, Anja Kunert for scientific advice, and Matthew Blango for thoughtful comments and discussions regarding this manuscript. We also acknowledge Antonia Lange for her experimental support in the isolation of environmental strains from 2016. G.P. would like to thank the Deutsche Forschungsgemeinschaft (DFG) CRC/Transregio 124 “Pathogenic fungi and their human host: Networks of interaction”, subproject INF & B5.

## AUTHOR CONTRIBUTIONS

O.K. and H.B.D. conceived the study. A.E.B. and J.B. collected samples and performed experiments. W.B. provided reagents and experimental advice. A.E.B., J.B., T.S.E., K.K, G.P., H.B.D., and O.K. analyzed the data and interpreted the results. A.E.B. wrote the manuscript with input from all authors.

## AUTHOR ORDER

For cases of equal authorship contribution, order was determined by alphabetical order.

---

**Supplemental Figure 1.**
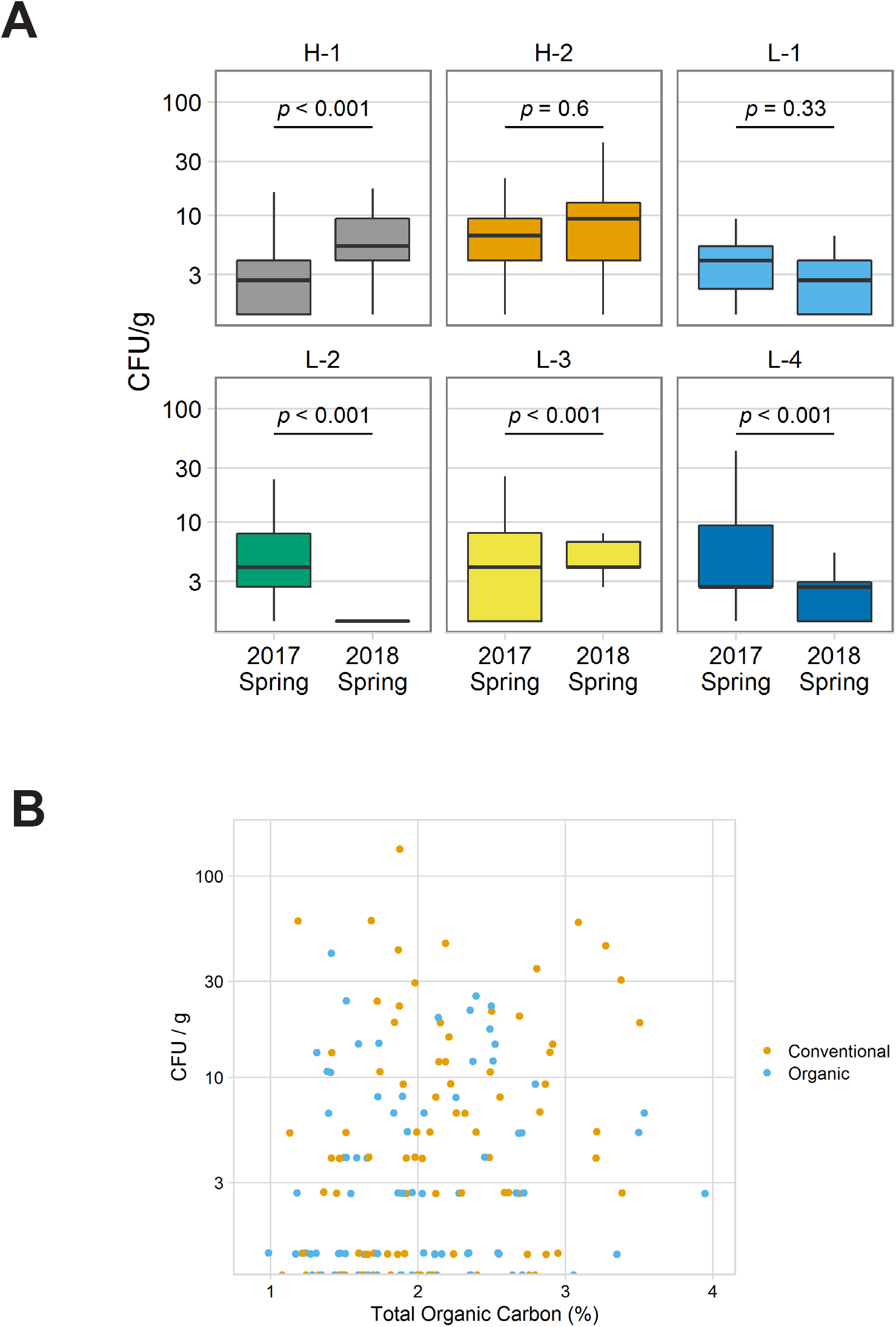
(A) Comparison of *A. fumigatus* abundance on apple fields sampled in spring 2017 and spring 2018. n = 50 soil samples per time point for fields H-1 and H-2; n = 25 soil samples per time point for fields L-1, L-2, L-3, L-4. *P* values calculated by Wilcoxon signed-rank test between the time points. (B) *A. fumigatus* abundance does not correlate with total organic carbon content of soil. n = 170 soil samples (10 per field from 17 fields on 10 different farms).

**Supplemental Figure 2.**
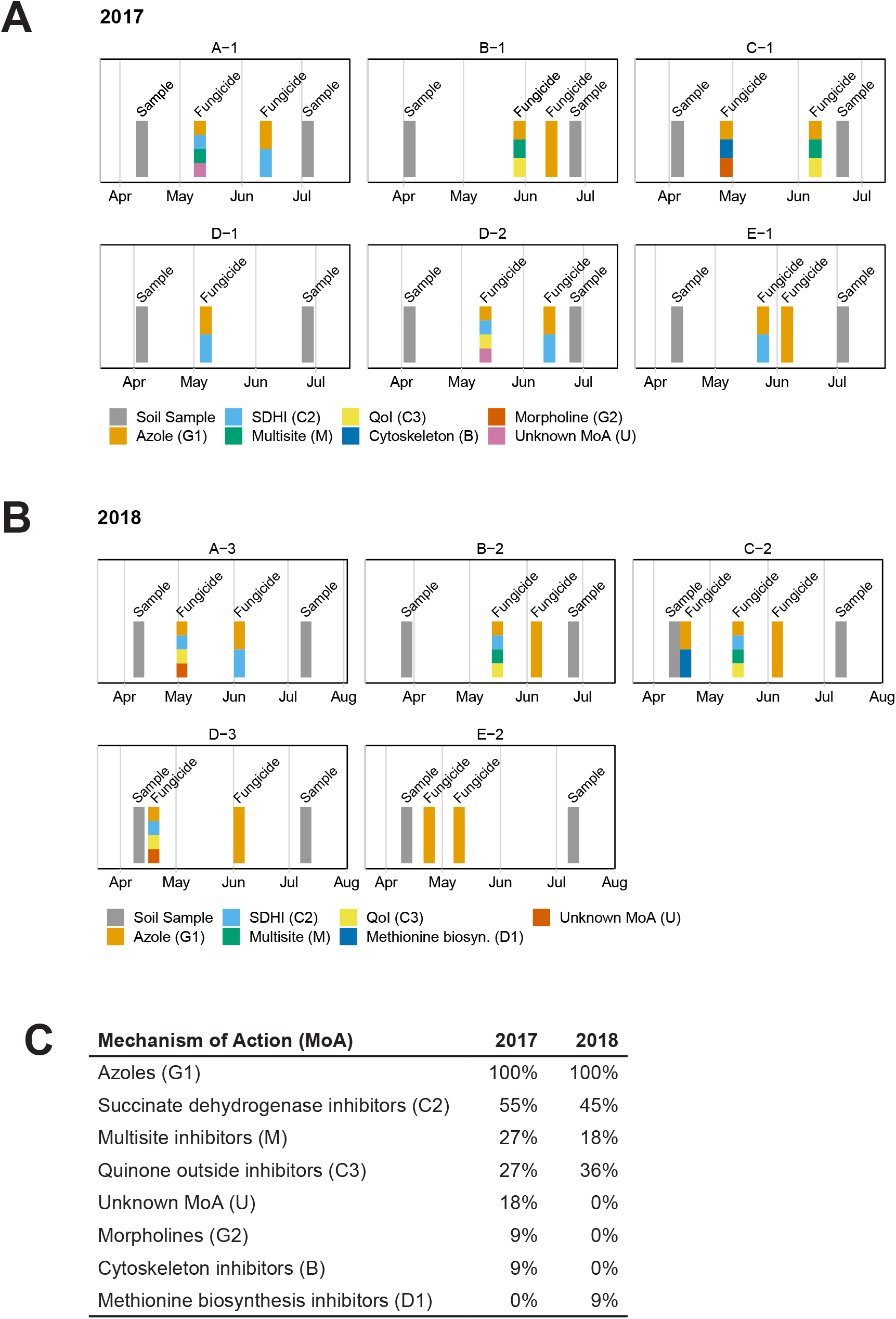
Fungicide history and schematic representation of the sampling time points and fungicide applications for the fields analyzed in this study. (A, B) Schematic representation of the sampling time points and fungicide applications for the fields sampled in 2017 (A) and 2018 (B). Soil sampling time points are marked in dark grey, while fungicide applications are noted in a combination of red and light grey indicating the relative proportion of azoles within the fungicide cocktail applied. Fungicide history was not available for fields on Farms H and L. (C) Overall frequency of fungicide classifications applied to the fields sampled during 2017 and 2018. Fungicides are categoried by their Fungicide Resistance Action Committee (FRAC) mechanism of action and a frequency of 100% means that a category was present in every fungicide application on every field analyzed that year.

**Supplemental Figure 3.**
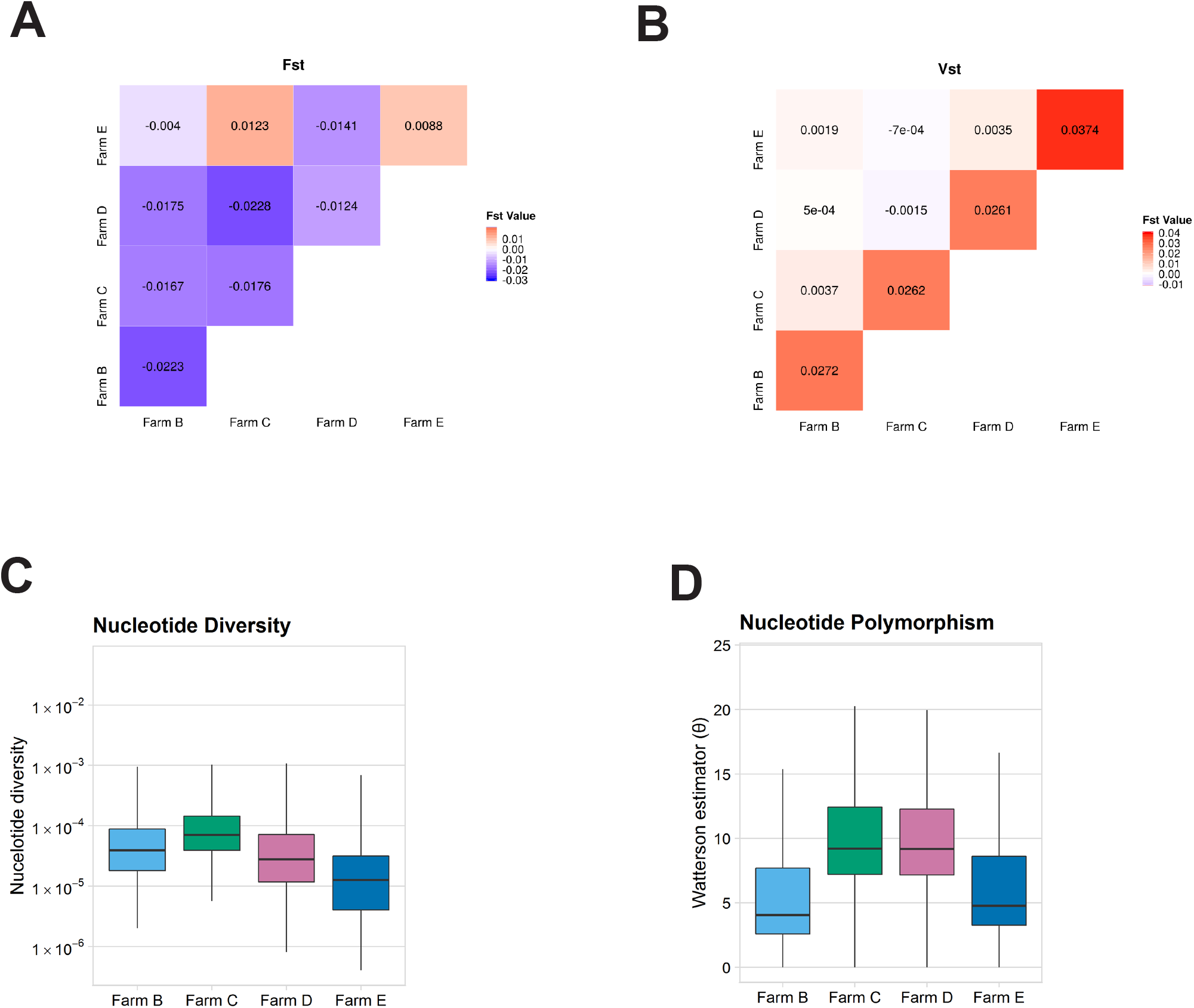
Population genetic of farms using isolates from before and after azole exposure. n = 16 isolates per farm. (A-B) Heat map representing weighted Weir and Cockerham F_ST_ values (A) and V_ST_ values (B) between farms. Blue colors represent more similarity between populations, while red represents greater population differentiation. (C-D) Comparison of nucleotide diversity (n) (C) and nucleotide polymor-phism (Watterson estimator or 8) (D) between farms. Values calculated from 5 kb sliding windows.

## REFERENCES

1. Leading International Fungal Education (LIFE): Invasive Aspergillosis. http://www.life-worldwide.org/fungal-diseases/invasive-aspergillosis.

2. Patterson TF, Iii RT, Denning DW, Fishman JA, Hadley S, Herbrecht R, Kontoyiannis DP, Marr KA, Morrison VA, Nguyen MH, Segal BH, Steinbach WJ, Stevens DA, Walsh TJ, Wingard JR. 2016. Practice Guidelines for the Diagnosis and Management of Aspergillosis : 2016 Update by the Infectious Diseases Society of America. Clinical Infectious Diseases doi:10.1093/cid/ciw326:1–60.

3. Buil JB, Snelders E, Denardi LB, Melchers WJG, Verweij PE. 2019. Trends in Azole Resistance in Aspergillus fumigatus, the Netherlands, 1994-2016. Emerg Infect Dis 25:176–178.

4. Steinmann J, Hamprecht A, Vehreschild MJ, Cornely OA, Buchheidt D, Spiess B, Koldehoff M, Buer J, Meis JF, Rath PM. 2015. Emergence of azole-resistant invasive aspergillosis in HSCT recipients in Germany. J Antimicrob Chemother 70:1522–6.

5. Chowdhary A, Kathuria S, Xu J, Meis JF. 2013. Emergence of Azole-Resistant Aspergillus fumigatus Strains due to Agricultural Azole Use Creates an Increasing Threat to Human Health. PLoS Pathogens 9:3–7.

6. Chowdhary A, Kathuria S, Xu J, Sharma C, Sundar G, Singh PK, Gaur SN, Hagen F, Klaassen CH, Meis JF. 2012. Clonal expansion and emergence of environmental multiple-triazole-resistant Aspergillus fumigatus strains carrying the TR(3)(4)/L98H mutations in the cyp51A gene in India. PLoS One 7:e52871.

7. Meis JF, Chowdhary A, Rhodes JL, Fisher MC, Verweij PE. 2016. Clinical implications of globally emerging azole resistance in Aspergillus fumigatus. Philos Trans R Soc Lond B Biol Sci 371.

8. Ballard E, Melchers WJG, Zoll J, Brown AJP, Verweij PE, Warris A. 2018. In-host microevolution of Aspergillus fumigatus: A phenotypic and genotypic analysis. Fungal Genet Biol 113:1–13.

9. Hare RK, Gertsen JB, Astvad KMT, Degn KB, Lokke A, Stegger M, Andersen PS, Kristensen L, Arendrup MC. 2019. In Vivo Selection of a Unique Tandem Repeat Mediated Azole Resistance Mechanism (TR120) in Aspergillus fumigatus cyp51A, Denmark. Emerg Infect Dis 25:577–580.

10. Berger S, El Chazli Y, Babu AF, Coste AT. 2017. Azole Resistance in Aspergillus fumigatus: A Consequence of Antifungal Use in Agriculture? Front Microbiol 8:1024.

11. Verweij PE, Snelders E, Kema GHJ, Mellado E, Melchers WJG. 2009. Azole resistance in Aspergillus fumigatus: a side-effect of environmental fungicide use? The Lancet Infectious Diseases 9:789–795.

12. Morton V, Staub T. 2008. A Short History of Fungicides. American Phytopathological Society Net Online Feature doi:American Phytopathological Society.

13. Maertens JA. 2004. History of the development of azole derivatives. Clin Microbiol Infect 10 Suppl 1:1–10.

14. Price CL, Parker JE, Warrilow AG, Kelly DE, Kelly SL. 2015. Azole fungicides - understanding resistance mechanisms in agricultural fungal pathogens. Pest Manag Sci 71:1054–8.

15. Anonymous. 2019. FRAC Code List 2019: Fungal control agents sorted by cross resistance pattern and mode of action. Fungicide Resistance Action Committee.

16. Fisher MC, Hawkins NJ, Sanglard D, Gurr SJ. 2018. Worldwide emergence of resistance to antifungal drugs challenges human health and food security. Science 360:739–742.

17. Mellado E, Garcia-Effron G, Alcazar-Fuoli L, Melchers WJ, Verweij PE, Cuenca-Estrella M, Rodriguez-Tudela JL. 2007. A new Aspergillus fumigatus resistance mechanism conferring in vitro cross-resistance to azole antifungals involves a combination of cyp51A alterations. Antimicrob Agents Chemother 51:1897–904.

18. Liu M, Zheng N, Li D, Zheng H, Zhang L, Ge H, Liu W. 2016. cyp51A-based mechanism of azole resistance in Aspergillus fumigatus: Illustration by a new 3D Structural Model of Aspergillus fumigatus CYP51A protein. Med Mycol 54:400–8.

19. Nash A, Rhodes J. 2018. Simulations of CYP51A from Aspergillus fumigatus in a model bilayer provide insights into triazole drug resistance. Med Mycol 56:361–373.

20. Zhang J, Snelders E, Zwaan BJ, Schoustra SE, Meis JF, van Dijk K, Hagen F, van der Beek MT, Kampinga GA, Zoll J, Melchers WJG, Verweij PE, Debets AJM. 2017. A Novel Environmental Azole Resistance Mutation in Aspergillus fumigatus and a Possible Role of Sexual Reproduction in Its Emergence. MBio 8.

21. Vermeulen E, Maertens J, Schoemans H, Lagrou K. 2012. Azole-resistant Aspergillus fumigatus due to TR46/Y121F/T289A mutation emerging in Belgium, July 2012. Euro Surveill 17.

22. Arendrup MC, Verweij PE, Mouton JW, Lagrou K, Meletiadis J. 2017. Multicentre validation of 4-well azole agar plates as a screening method for detection of clinically relevant azole-resistant Aspergillus fumigatus. J Antimicrob Chemother 72:3325–3333.

23. Buil JB, van der Lee HAL, Rijs A, Zoll J, Hovestadt J, Melchers WJG, Verweij PE. 2017. Single-Center Evaluation of an Agar-Based Screening for Azole Resistance in Aspergillus fumigatus by Using VIPcheck. Antimicrob Agents Chemother 61.

24. Snelders E, Camps SMT, Karawajczyk A, Schaftenaar G, Kema GHJ, van der Lee HA, Klaassen CH, Melchers WJG, Verweij PE. 2012. Triazole fungicides can induce cross-resistance to medical triazoles in Aspergillus fumigatus. PLoS ONE 7.

25. Garcia-Rubio R, Monzon S, Alcazar-Fuoli L, Cuesta I, Mellado E. 2018. Genome-Wide Comparative Analysis of Aspergillus fumigatus Strains: The Reference Genome as a Matter of Concern. Genes (Basel) 9.

26. Bowyer P, Denning DW. 2014. Environmental fungicides and triazole resistance in Aspergillus. Pest Management Science 70:173–178.

27. Rhodes J. 2019. Rapid Worldwide Emergence of Pathogenic Fungi. Cell Host Microbe 26:12–14.

28. Resendiz Sharpe A, Lagrou K, Meis JF, Chowdhary A, Lockhart SR, Verweij PE, group IEARSw. 2018. Triazole resistance surveillance in Aspergillus fumigatus. Med Mycol 56:83–92.

29. Bader O, Weig M, Reichard U, Lugert R, Kuhns M, Christner M, Held J, Peter S, Schumacher U, Buchheidt D, Tintelnot K, Gross U, MykoLabNet DP. 2013. cyp51A-Based mechanisms of Aspergillus fumigatus azole drug resistance present in clinical samples from Germany. Antimicrob Agents Chemother 57:3513–7.

30. Bader O, Tünnermann J, Dudakova A, Tangwattanachuleeporn M, Weig M, Groß U, Hoberg N, Geibel S, Vogel E, Büntzel J, Springer J, Lehning LY, Schädel C, Antweiler E, Metzger L, Zautner A, Buchheidt D, Spiess B, Hamprecht A, Steinmann J, Rößler S, Wiegmann S, Klingebiel S, Loock AC, Hegewald J, Hassenpflug M, Aurin A, Szymczak J, Diffloth N, Kuhns M. 2015. Environmental isolates of azole-resistant Aspergillus fumigatus in Germany. Antimicrobial Agents and Chemotherapy 59:4356–4359.

31. Alvarez-Moreno C, Lavergne RA, Hagen F, Morio F, Meis JF, Le Pape P. 2019. Fungicide-driven alterations in azole-resistant Aspergillus fumigatus are related to vegetable crops in Colombia, South America. Mycologia 111:217–224.

32. Prigitano A, Venier V, Cogliati M, De Lorenzis G, Esposto MC, Tortorano AM. 2014. Azole-resistant aspergillus fumigatus in the environment of Northern Italy, May 2011 to June 2012. Eurosurveillance 19:1–7.

33. Mortensen KL, Mellado E, Lass-Fl??rl C, Rodriguez-Tudela JL, Johansen HK, Arendrup MC. 2010. Environmental study of azole-resistant Aspergillus fumigatus and other aspergilli in Austria, Denmark, and Spain. Antimicrobial Agents and Chemotherapy 54:4545–4549.

34. Sewell TR, Zhang Y, Brackin AP, Shelton JMG, Rhodes J, Fisher MC. 2019. Elevated prevalence of azole resistant Aspergillus fumigatus in urban versus rural environments in the United Kingdom. Antimicrob Agents Chemother doi:10.1128/AAC.00548-19.

35. Jeanvoine A, Rocchi S, Reboux G, Crini N, Crini G, Millon L. 2017. Azole-resistant Aspergillus fumigatus in sawmills of Eastern France. J Appl Microbiol doi:10.1111/jam.13488.

36. Cox KD, Bryson PK, Schnabel G. 2007. Instability of Propiconazole Resistance and Fitness in Monilinia fructicola. Phytopathology 97:448–53.

37. Calo S, Shertz-Wall C, Lee SC, Bastidas RJ, Nicolas FE, Granek JA, Mieczkowski P, Torres-Martinez S, Ruiz-Vazquez RM, Cardenas ME, Heitman J. 2014. Antifungal drug resistance evoked via RNAi-dependent epimutations. Nature 513:555–8.

38. Schoustra SE, Debets AJM, Rijs A, Zhang J, Snelders E, Leendertse PC, Melchers WJG, Rietveld AG, Zwaan BJ, Verweij PE. 2019. Environmental Hotspots for Azole Resistance Selection of Aspergillus fumigatus, the Netherlands. Emerg Infect Dis 25:1347–1353.

39. Abdolrasouli A, Rhodes J, Beale MA, Hagen F, Rogers TR, Chowdhary A, Meis JF, Armstrong-James D, Fisher MC. 2015. Genomic Context of Azole Resistance Mutations in Aspergillus fumigatus Determined Using Whole-Genome Sequencing. MBio 6:e00536.

40. Dunne K, Hagen F, Pomeroy N, Meis JF, Rogers TR. 2017. Intercountry Transfer of Triazole-Resistant Aspergillus fumigatus on Plant Bulbs. Clin Infect Dis 65:147–149.

41. Alvarez-Moreno C, Lavergne RA, Hagen F, Morio F, Meis JF, Le Pape P. 2017. Azole-resistant Aspergillus fumigatus harboring TR34/L98H, TR46/Y121F/T289A and TR53 mutations related to flower fields in Colombia. Sci Rep 7:45631.

42. Guinea J, Verweij PE, Meletiadis J, Mouton JW, Barchiesi F, Arendrup MC, Subcommittee on Antifungal Susceptibility Testing of the EECfAST. 2018. How to: EUCAST recommendations on the screening procedure E.Def 10.1 for the detection of azole resistance in Aspergillus fumigatus isolates using four-well azole-containing agar plates. Clin Microbiol Infect doi:10.1016/j.cmi.2018.09.008.

43. Basenko EY, Pulman JA, Shanmugasundram A, Harb OS, Crouch K, Starns D, Warrenfeltz S, Aurrecoechea C, Stoeckert CJ, Jr., Kissinger JC, Roos DS, Hertz-Fowler C. 2018. FungiDB: An Integrated Bioinformatic Resource for Fungi and Oomycetes. J Fungi (Basel) 4.

44. Li H, Durbin R. 2009. Fast and accurate short read alignment with Burrows-Wheeler transform. Bioinformatics 25:1754–60.

45. McKenna A, Hanna M, Banks E, Sivachenko A, Cibulskis K, Kernytsky A, Garimella K, Altshuler D, Gabriel S, Daly M, DePristo MA. 2010. The Genome Analysis Toolkit: a MapReduce framework for analyzing next-generation DNA sequencing data. Genome Res 20:1297–303.

46. Boeva V, Popova T, Bleakley K, Chiche P, Cappo J, Schleiermacher G, Janoueix-Lerosey I, Delattre O, Barillot E. 2012. Control-FREEC: a tool for assessing copy number and allelic content using next-generation sequencing data. Bioinformatics 28:423–5.

47. Danecek P, Auton A, Abecasis G, Albers CA, Banks E, DePristo MA, Handsaker RE, Lunter G, Marth GT, Sherry ST, McVean G, Durbin R, Genomes Project Analysis G. 2011. The variant call format and VCFtools. Bioinformatics 27:2156–8.

48. Edgar RC. 2004. MUSCLE: multiple sequence alignment with high accuracy and high throughput. Nucleic Acids Res 32:1792–7.

49. Price MN, Dehal PS, Arkin AP. 2010. FastTree 2--approximately maximum-likelihood trees for large alignments. PLoS One 5:e9490.

50. Letunic I, Bork P. 2019. Interactive Tree Of Life (iTOL) v4: recent updates and new developments. Nucleic Acids Res 47:W256–W259.

51. Korneliussen TS, Moltke I, Albrechtsen A, Nielsen R. 2013. Calculation of Tajima's D and other neutrality test statistics from low depth next-generation sequencing data. BMC Bioinformatics 14:289.

52. Redon R, Ishikawa S, Fitch KR, Feuk L, Perry GH, Andrews TD, Fiegler H, Shapero MH, Carson AR, Chen W, Cho EK, Dallaire S, Freeman JL, Gonzalez JR, Gratacos M, Huang J, Kalaitzopoulos D, Komura D, MacDonald JR, Marshall CR, Mei R, Montgomery L, Nishimura K, Okamura K, Shen F, Somerville MJ, Tchinda J, Valsesia A, Woodwark C, Yang F, Zhang J, Zerjal T, Zhang J, Armengol L, Conrad DF, Estivill X, Tyler-Smith C, Carter NP, Aburatani H, Lee C, Jones KW, Scherer SW, Hurles ME. 2006. Global variation in copy number in the human genome. Nature 444:444–54.

