## Supplemental Materials & Methods, Supplemental Tables 1-5 for "Effects of agricultural fungicide use on *Aspergillus fumigatus* abundance, antifungal susceptibility, and population structure"

### SUPPLEMENTAL MATERIALS & METHODS

#### Additional information on field samling and the naming of fields

Six of the sampled farms practiced biological (organic) measures of land management without azole application, while seven practiced integrated pest management, including application of azole fungicides. To differentiate the year the sampling occurred, a corresponding suffix was added to the Field ID (*e.g.* D-2-17 identifies the second field sampled on farm D in the year 2017). Due to crop rotation, the same field could not be surveyed over subsequent years, nor was it possible to always sample the same number of fields from each site each year. To avoid confusion, when we returned to farms the following year, fields were assigned new numbers starting with where the labeling system left off the previous year. (*e.g.* on farm D, two field were sampled in 2017, so in 2018 the labeling system started with D-3). Fields that were sampled over subsequent years, which was only possible for apple orchards, maintained the same numbering system (*e.g.* L-4-17 and L-4-18 are the same fields). In the case where two fields of the same agricultural type on the same farm were sampled, the number of soil samples was divided between the two fields and 25 samples per field were collected.

#### Assessment of TOC content

Soil samples were air dried and sieved through a 2 mm stainless steel mesh to remove larger organic particles and stones. Samples were then manually ground to a fine powder and divided into two parts. The first fraction (approximately 1000 mg) was used to measure total carbon (TC) content by total combustion with oxygen at 1000°C using an elemental macro-analyzer (Elementar Vario Max Cube, Langenselbold, Germany). The second fraction (appr. 200 mg) was used to determine the total inorganic fraction (TIC) present by volumetrically adding hydrochloric acid and recording the volume of $CO_2$ produced (also using an Elementar Vario Max Cube, Langenselbold, Germany). The final total organic carbon (TOC) content in the soil was calculated by subtracting the total carbon (TC) from the total inorganic carbon (TIC).

**Supplemental Table 1**. Summary of the agricultural sites surveyed 2016- 2018.

| Farm | Type of Agriculture | Crop | Soil samples (n) | | | *A. fumigatus* isolates (n) | | |
|---|---|---|---|---|---|---|---|---|
| | | | 2016 | 2017 | 2018 | 2016 | 2017 | 2018 |
| A | Conventional and organic | Cereal | 150 | 150 | 150 | 232 | 94 | 81 |
| B | Conventional | Cereal | 100 | 100 | 100 | 114 | 266 | 335 |
| C | Conventional | Cereal | 100 | 100 | 100 | 29 | 123 | 37 |
| D | Conventional | Cereal | 100 | 100 | 100 | 65 | 228 | 287 |
| E | Conventional | Cereal | 100 | 100 | 100 | 61 | 131 | 165 |
| F | Organic | Cereal | 50 | 50 | 25 | 73 | 87 | 15 |
| G | Organic | Cereal | 50 | 50 | 50 | 29 | 87 | 13 |
| H | Conventional and organic | Apple | 100 | 150 | 150 | 158 | 301 | 421 |
| K | Organic | Cereal | 50 | 50 | 50 | 45 | 26 | 6 |
| L | Conventional and organic | Apple | 100 | 150 | 150 | 91 | 184 | 162 |
| | | Total | 900 | 1000 | 975 | 897 | 1527 | 1522 |

**Supplemental Table 2**. Resistance summary of the fields sampled during the spring of 2017 (A) and 2018 (B).

**(A)**

| Type of agriculture | Field | Crop | n tested | Proportion of isolates that grow at the concentration indicated<br>DIF (1 mg/L) | TEB (2 mg/L) | ITR (4 mg/L) | VOR (2 mg/L) | POS (0.5 mg/L) |
|---|---|---|---|---|---|---|---|---|
| Conventional | A-1-17 | Cereal | 20 | 0.15 | 0.00 | 0.00 | 0.00 | 0.00 |
| | B-1-17 | Cereal | 20 | 0.25 | 0.05 | 0.00 | 0.00 | 0.00 |
| | C-1-17 | Cereal | 20 | 0.10 | 0.00 | 0.00 | 0.00 | 0.00 |
| | D-1-17 | Cereal | 20 | 0.25 | 0.00 | 0.00 | 0.00 | 0.00 |
| | D-2-17 | Cereal | 20 | 0.40 | 0.05 | 0.00 | 0.00 | 0.00 |
| | E-1-17 | Cereal | 20 | 0.15 | 0.05 | 0.00 | 0.00 | 0.00 |
| | H-1-17 | Apple | 20 | 0.15 | 0.05 | 0.00 | 0.00 | 0.00 |
| | L-1-17 | Apple | 20 | 0.15 | 0.00 | 0.00 | 0.00 | 0.00 |
| | L-4-17 | Apple | 20 | 0.55 | 0.00 | 0.00 | 0.00 | 0.00 |
| Organic | A-2-17 | Cereal | 20 | 0.35 | 0.15 | 0.05 | 0.00 | 0.00 |
| | F-1-17 | Cereal | 20 | 0.25 | 0.00 | 0.00 | 0.00 | 0.00 |
| | G-1-17 | Cereal | 20 | 0.25 | 0.10 | 0.05 | 0.00 | 0.00 |
| | G-2-17 | Cereal | 20 | 0.15 | 0.00 | 0.00 | 0.00 | 0.00 |
| | K-1-17 | Cereal | 20 | 0.25 | 0.00 | 0.00 | 0.00 | 0.00 |
| | H-2-17 | Apple | 20 | 0.25 | 0.00 | 0.00 | 0.00 | 0.00 |
| | L-2-17 | Apple | 20 | 0.10 | 0.05 | 0.00 | 0.00 | 0.00 |
| | L-3-17 | Apple | 20 | 0.35 | 0.25 | 0.00 | 0.00 | 0.00 |

**(B)**

| Type of agriculture | Field | Crop | n tested | Proportion of isolates that grow at the concentration indicated<br>DIF (1 mg/L) | TEB (2 mg/L) | ITR (4 mg/L) | VOR (2 mg/L) | POS (0.5 mg/L) |
|---|---|---|---|---|---|---|---|---|
| Conventional | A-3-18 | Cereal | 11 | 0.27 | 0.00 | 0.00 | 0.00 | 0.00 |
| | B-2-18 | Cereal | 20 | 0.10 | 0.05 | 0.00 | 0.00 | 0.00 |
| | C-2-18 | Cereal | 11 | 0.09 | 0.00 | 0.00 | 0.00 | 0.00 |
| | D-3-18 | Cereal | 20 | 0.10 | 0.05 | 0.00 | 0.00 | 0.00 |
| | E-2-18 | Cereal | 20 | 0.10 | 0.10 | 0.00 | 0.00 | 0.00 |
| | H-1-18 | Apple | 20 | 0.20 | 0.10 | 0.00 | 0.00 | 0.00 |
| | L-1-18 | Apple | 18 | 0.11 | 0.00 | 0.00 | 0.00 | 0.00 |
| | L-4-18 | Apple | 12 | 0.17 | 0.00 | 0.00 | 0.00 | 0.00 |
| Organic | A-4-18 | Cereal | 20 | 0.40 | 0.20 | 0.00 | 0.00 | 0.00 |
| | F-2-18 | Cereal | 10 | 0.40 | 0.00 | 0.00 | 0.00 | 0.00 |
| | G-3-18 | Cereal | 12 | 0.50 | 0.00 | 0.00 | 0.00 | 0.00 |
| | K-2-18 | Cereal | 5 | 0.00 | 0.00 | 0.00 | 0.00 | 0.00 |
| | H-2-18 | Apple | 20 | 0.20 | 0.05 | 0.00 | 0.00 | 0.00 |
| | L-2-18 | Apple | 6 | 0.17 | 0.00 | 0.00 | 0.00 | 0.00 |
| | L-3-18 | Apple | 8 | 0.38 | 0.00 | 0.00 | 0.00 | 0.00 |

**Supplemental Table 3**. Resistance summary of the fields sampled before and after the vegetative period and azole application in 2017 (A) and 2018 (B).

**(A)**

| | | | | Proportion of isolates that grow at the concentration indicated | | | | |
|---|---|---|---|---|---|---|---|---|
| Field | Crop | Time Period | n tested | DIF (1 mg/L) | TEB (2 mg/L) | ITR (4 mg/L) | VOR (2 mg/L) | POS (0.5 mg/L) |
| B-1-17 | Cereal | Before | 20 | 0.25 | 0.05 | 0.00 | 0.00 | 0.00 |
| | | After | 20 | 0.50 | 0.15 | 0.00 | 0.00 | 0.00 |
| C-1-17 | Cereal | Before | 20 | 0.10 | 0.00 | 0.00 | 0.00 | 0.00 |
| | | After | 20 | 0.45 | 0.15 | 0.10 | 0.00 | 0.00 |
| D-1-17 | Cereal | Before | 20 | 0.25 | 0.00 | 0.00 | 0.00 | 0.00 |
| | | After | 20 | 0.50 | 0.25 | 0.00 | 0.00 | 0.00 |
| D-2-17 | Cereal | Before | 20 | 0.40 | 0.05 | 0.00 | 0.00 | 0.00 |
| | | After | 20 | 0.35 | 0.05 | 0.00 | 0.00 | 0.00 |
| E-1-17 | Cereal | Before | 20 | 0.15 | 0.05 | 0.00 | 0.00 | 0.00 |
| | | After | 15 | 0.47 | 0.07 | 0.00 | 0.00 | 0.00 |
| H-1-17 | Apple | Before | 20 | 0.15 | 0.05 | 0.00 | 0.00 | 0.00 |
| | | After | 20 | 0.55 | 0.20 | 0.00 | 0.00 | 0.00 |
| L-4-17 | Apple | Before | 20 | 0.55 | 0.00 | 0.00 | 0.00 | 0.00 |
| | | After | 20 | 0.50 | 0.10 | 0.00 | 0.00 | 0.00 |

**(B)**

| | | | | Proportion of isolates that grow at the concentration indicated | | | | |
|---|---|---|---|---|---|---|---|---|
| Field | Crop | Time Period | n tested | DIF (1 mg/L) | TEB (2 mg/L) | ITR (4 mg/L) | VOR (2 mg/L) | POS (0.5 mg/L) |
| A-3-18 | Cereal | Before | 11 | 0.27 | 0.00 | 0.00 | 0.00 | 0.00 |
| | | After | 11 | 0.27 | 0.18 | 0.00 | 0.00 | 0.00 |
| B-2-18 | Cereal | Before | 20 | 0.10 | 0.05 | 0.00 | 0.00 | 0.00 |
| | | After | 20 | 0.25 | 0.10 | 0.05 | 0.00 | 0.00 |
| C-2-18 | Cereal | Before | 11 | 0.09 | 0.00 | 0.00 | 0.00 | 0.00 |
| | | After | 16 | 0.25 | 0.06 | 0.00 | 0.00 | 0.00 |
| D-3-18 | Cereal | Before | 20 | 0.10 | 0.05 | 0.00 | 0.00 | 0.00 |
| | | After | 20 | 0.35 | 0.10 | 0.05 | 0.00 | 0.00 |
| H-1-18 | Apple | Before | 20 | 0.20 | 0.10 | 0.00 | 0.00 | 0.00 |
| | | After | 20 | 0.50 | 0.10 | 0.00 | 0.00 | 0.00 |
| L-1-18 | Apple | Before | 18 | 0.11 | 0.00 | 0.00 | 0.00 | 0.00 |
| | | After | 10 | 0.50 | 0.00 | 0.00 | 0.00 | 0.00 |
| L-4-18 | Apple | Before | 12 | 0.17 | 0.00 | 0.00 | 0.00 | 0.00 |
| | | After | 12 | 0.58 | 0.08 | 0.00 | 0.00 | 0.00 |

**Supplemental Table 4**. MICs for medical and agricultural azoles in isolates displaying resistance to at least one medical azole.

| Strain | *cyp51a* genotype[+] | ITR | POS | VOR | DIF | TEB | AMB |
|---|---|---|---|---|---|---|---|
| 2016-072 | TR34/L98H | >8 | 1 | 4 | >8 | >8 | 0.25 |
| 2016-262 | TR34/L98H | >8 | 1 | 4 | >8 | >8 | 0.25 |
| 2016-334 | TR34/L98H | >8 | 0.5 | 2 | 8 | >8 | 0.25 |
| 2016-375 | TR34/L98H | >8 | 0.5 | 4 | 4 | >8 | 0.5 |
| 2016-439 | TR34/L98H | >8 | 1 | 4 | >8 | >8 | 0.25 |
| 2016-698 | TR34/L98H | >8 | 1 | 4 | >8 | >8 | 0.25 |
| 2017-105 | TR34/L98H | >8 | 1 | 4 | >8 | >8 | 0.25 |
| 2017-402 | TR34/L98H | >8 | 1 | 2 | 8 | >8 | 0.25 |
| 2017-415 | TR34/L98H | >8 | 1 | 4 | 8 | >8 | 0.25 |
| 2016-106 | WT | >8 | 0.25 | 1 | 4 | 8 | 0.5 |
| 2017-153 | WT | >8 | 0.5 | 2 | 4 | 8 | 0.25 |
| 2016-684 | WT | 8 | 0.5 | 1 | 4 | 8 | 0.5 |
| 2016-313 | WT | 4 | 0.25 | 1 | 8 | >8 | 0.5 |
| 2016-675 | WT | 4 | 0.5 | 2 | 8 | 8 | 0.5 |
| 2016-369 | WT | 2 | 0.125 | 1 | 2 | 4 | 0.25 |
| 2018-214 | WT | 2 | 0.25 | 1 | 4 | 4 | 0.25 |
| 2018-272 | WT | 2 | 0.125 | 1 | 4 | 4 | 0.25 |

Abbreviations: amphotericin B = AMB; difenoconazole = DIF; itraconazole = ITR; posaconazole = POS; tebuconazole = TEB; voriconazole = VOR; WT = wild type.
[+]Relative to ATCC36607.

**Supplemental Table 5**. Primers used for amplifying and sequencing *cyp51a*.

| Primer | Sequence | Use |
|---|---|---|
| cyp51a_F | 5'-CGTAGCAAGGGAGAAGGAAA | Amplification and sequencing |
| cyp51a_R | 5'-CACCTATTCCGATCACACCA | Amplification and sequencing |
| cyp51a_internal | 5'-ATGTCAATGCGGAAGAGGTC | Sequencing |
